# Microglia regulate developmental transitions in synaptic plasticity rules in the medial prefrontal cortex

**DOI:** 10.1101/2025.06.16.659241

**Authors:** Mio Tajiri, Hidefumi Omi, Airi Inoue, Hayato Etani, Takeshi Sawada, Yusuke Iino, Sho Yagishita

**Affiliations:** Laboratory of Structural Physiology, Center for Disease Biology and Integrative Medicine, Graduate School of Medicine, The University of Tokyo, Tokyo, Japan; International Research Center for Neurointelligence (WPI-IRCN), UTIAS, The University of Tokyo, Tokyo, Japan

## Abstract

Synaptic plasticity is governed by rules that determine how patterns of neural activity are converted into changes in synaptic weights. While neocortical plasticity rules appear to shift from Hebbian to three-factor forms during development, the cellular mechanisms underlying this transition remain unclear. Here, we show that microglia mediate the developmental transition of plasticity rules at individual dendritic spines in the medial prefrontal cortex (mPFC) during juvenile-to-adolescent maturation. After maturation, microglia suppressed Hebbian activity–induced spine enlargement via a humoral factor. This suppression was relieved by noradrenaline through microglia-intrinsic cAMP signaling, thereby enabling a three-factor rule for gating plasticity. The adolescent mPFC required three-factor plasticity signaling for the acquisition of socially learned fear. Local microglial ablation enhanced learning efficiency but also altered pain sensitivity in a learning-dependent manner, suggesting that microglial suppression safeguards pre-existing emotional circuits. Together, these findings identify microglia as critical regulators of developmental plasticity-rule transitions in the mPFC.

---

Synaptic plasticity is the fundamental neural mechanism for learning (1, 2), which is controlled by “rules” that determine how patterns of neural activity are converted into changes in synaptic weights (3–10). In the neocortex, these rules appear to shift across maturation (5, 11–14). During childhood, neocortical circuits can remodel according to the Hebbian rule, strengthening synapses through coincident pre- and postsynaptic activity driven by glutamate (5, 15–18). From adolescence onward, however, Hebbian weight updates become less effective (14, 18–22), and plasticity increasingly depends on neuromodulatory contexts involving reward or punishment (23–29). The requirement for neuromodulators such as monoamines as a third factor, in addition to the two factors of pre- and postsynaptic activity, is known as the three-factor rule (7–9). This rule enables neural circuits to selectively update synaptic weights based on behaviorally relevant feedback. As excessive plasticity can destabilize established circuits (30, 31), this shift may stabilize existing representations while permitting updates only in select contexts, including emotionally salient ones, thereby contributing to the neocortical capacity for long-term information storage (30–33). Although this framework highlights both reduced Hebbian plasticity and increased neuromodulator-dependent mechanisms as central to the shift in synaptic rules in the neocortex, the exact cellular mechanisms are not well understood.

At excitatory glutamatergic synapses, dendritic spines function as the basic units for synaptic weight updating (34, 35). In the neocortex, spine volume reflects AMPA receptor content (36, 37) and learning induces input-specific structural long-term potentiation (sLTP) of dendritic spines (38–40). In *ex vivo* studies using hippocampal slices, the Hebbian pairing of glutamatergic input with postsynaptic spikes reliably induces sLTP (41–43). In contrast, glutamatergic stimulation induces far less prominent sLTP in the mature neocortex (44), suggesting the necessity of a three-factor rule. Consistent with this idea, neuromodulators such as noradrenaline (NA) enhance spine plasticity in vivo (45), supporting a role for neuromodulatory reinforcement in sLTP alongside concurrent glutamatergic synaptic activity during learning (38–40). Despite extensive work on the role of sLTP in neocortical learning, it remains unclear how Hebbian pairing-induced sLTP is suppressed in the mature neocortex and how this suppression interacts with neuromodulator-driven facilitation.

Both neuron-intrinsic maturation and glial mechanisms have been proposed to developmentally regulate plasticity, potentially contributing to Hebbian plasticity suppression. Neurons may raise the threshold for glutamatergic induction of plasticity (12, 18, 46), making neuromodulatory input increasingly necessary. More recent studies highlight glial roles in synaptic regulation. In particular, microglia regulate synapse number and spine turnover during developmental maturation (47–50) and respond to noradrenergic signals (51, 52) to influence synaptic function (52). However, it remains unclear whether microglia actively participate in defining rules for weight updates at the moment neural activity occurs, or instead modulate synaptic changes hours later, given the limited temporal resolution of current *in vivo* studies. Furthermore, conflicting findings report that microglia can either enhance or restrict day-scale structural remodeling (47, 49, 52–57), leaving their contribution to plasticity rules unresolved. Within the neocortex, the medial prefrontal cortex (mPFC), a key region for neuromodulator-dependent emotional learning (26, 58–62), undergoes synaptic maturation during adolescence with microglial involvement (63). Thus, the mPFC provides a critical site to investigate whether microglia contribute to the adolescent shift from Hebbian to three-factor learning rules, thereby helping establish emotional learning while maintaining pre-existing circuits.

In this study, we paired two-photon glutamate uncaging with action potential firing to impose a Hebbian pairing activity pattern on individual dendritic spines while simultaneously visualizing those spines and nearby microglia in acute mPFC slices. We observed a developmental decline in Hebbian pairing-induced sLTP and identified microglia as key mediators of this suppression, which depended on humoral signaling rather than process surveillance. We then tested whether sLTP could be induced by a three-factor rule by introducing neuromodulatory signals of NA, and found that NA relieved microglial suppression to gate sLTP through microglia-intrinsic signaling pathways. To examine the behavioral relevance of this plasticity mechanism, we used social fear learning (62, 64–68), which is rapidly acquired within a day and involves mPFC remodeling during learning (69). Finally, we asked whether microglia-mediated suppression of plasticity contributes to maintaining the functional integrity of emotional circuits during learning.

## Microglial suppression of spine enlargement during development

We first characterized neocortical sLTP induced by the Hebbian pattern of coincident pre- and postsynaptic activity in adolescent and juvenile mice, both in the presence and absence of microglia. To ablate microglia, we locally injected clodronate into the mPFC and confirmed nearly 90% ablation across a broad region of the mPFC 1–2 days after injection (Supplementary Figure 1A−C) (53). To visualize microglia in acute slices, we generated double-transgenic mice carrying a microglia-specific Cre driver allele, Tmem119-CreERT2, which preserves endogenous transmembrane protein 119 (Tmem119) gene expression through a P2A sequence (70), along with a Cre-dependent tdTomato reporter allele (Ai14), achieving more than 99% labeling in the mPFC (Figure 1A, Supplementary Figure 1D−G).

**Figure 1.**
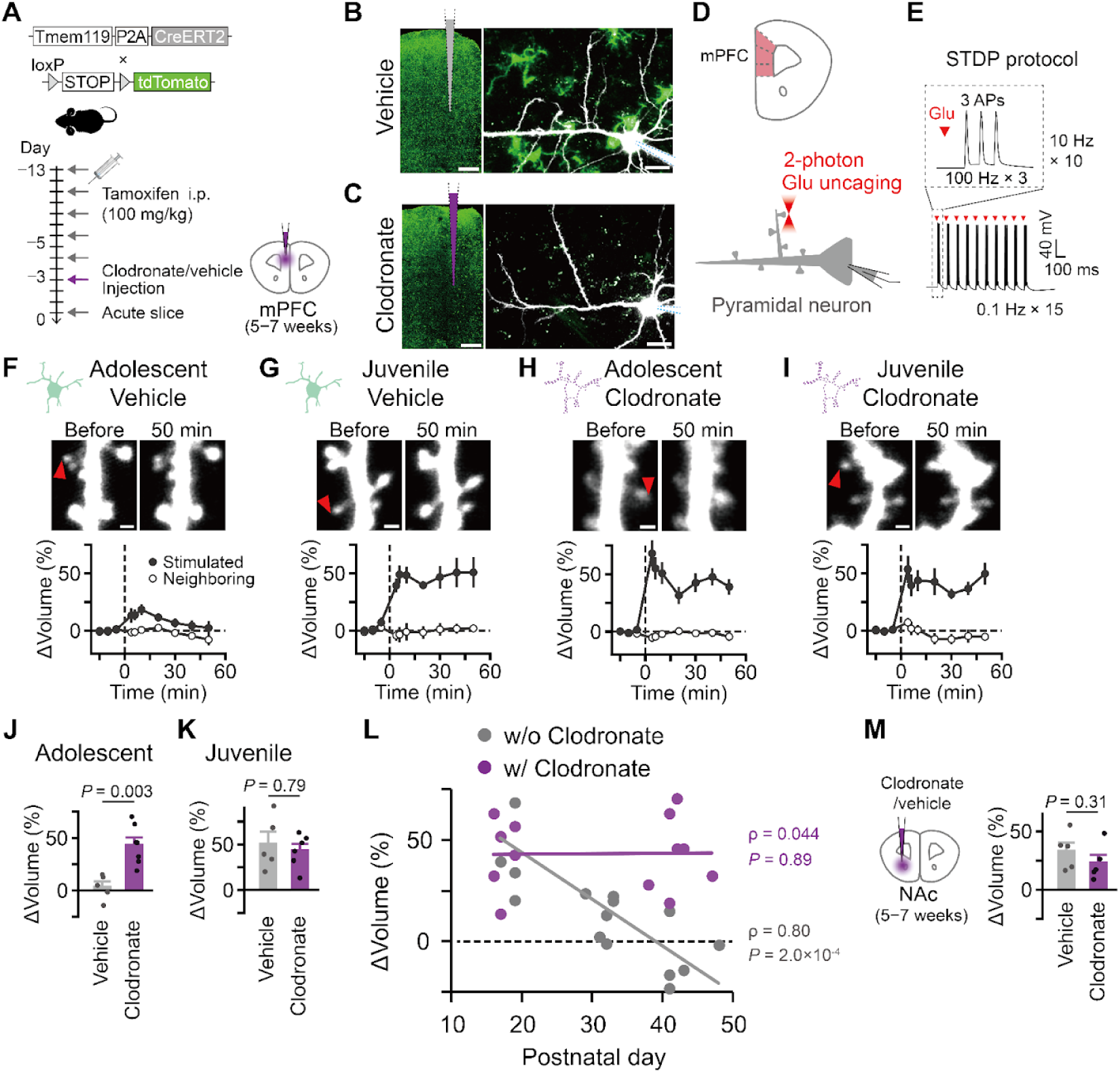
Microglial suppression of spine enlargement. **A,** Schematic of a typical experimental timeline in adolescent (5–7 weeks old) transgenic mice. **B,C,** Representative images of fixed slices with confocal microscopy (left) and acute slices with 2-photon microscopy (right) after vehicle (B) or clodronate (C) injections. Fluorescences of tdTomato for microglia (green) and Alexa 488 for neurons (white) are shown. Dotted lines indicate injection traces. Scale bars, 500 μm (left) and 20 μm (right). **D,** Schematic of a brain slice containing the mPFC (top) and the experimental configuration for two-photon uncaging on dendritic spines of a pyramidal neuron (bottom). Caged-glutamate (CDNI-Glutamate, 4 mM) was applied using a puff pipette. Two-photon laser was directed to the tip of the spine head (720nm, 0.6 ms, 4−6 mW, adjusted depending on the depth of the dendrite). Small 3−4 spines located on the first to third branches of the apical dendrites (20 μm to 50 μm below the surface of the slice) were selected for plasticity induction. Glu, glutamate. **E,** Schematic of the spike-timing-dependent plasticity (STDP) protocol. Each train consisted of 10 bursts (3 action potentials [APs] at 100 Hz) delivered at 10 Hz, where glutamate uncaging was paired with action potential within 10 ms. The train was repeated 15 times at 0.1 Hz. **F–I,** Representative images of dendritic spines before and 50 minutes after STDP stimulation (top) and time courses of spine volume changes (bottom) in juvenile mice (2−3 weeks old) (F,H) and the adolescent mice (G, I) after vehicle (F, G) or clodronate (H, I) injections. Red arrows indicate stimulated spines. Scale bars, 1 μm. **J,K,** Plots of spine volume changes. Volume changes were averaged over 40−50 minutes. **L,** Plots of spine volume changes across ages. Statistical analysis was performed using Spearman’s rank correlation coefficient test. **M,** Schematic of clodronate injection into the nucleus accumbens (NAc) in adolescent mice (5–7 weeks old) (left) and plots of spine volume changes (right). Error bars represent S.E.M. Mann–Whitney U test used for statistical analysis in (J,K,M).

Coronal acute slices containing the mPFC were prepared from adolescent (5 to 7 weeks) and juvenile (2 to 3 weeks of age) mice. Layer 5 pyramidal neurons in the mPFC were subjected to whole-cell recordings and perfused with Alexa 488 to visualize dendritic spines using two-photon microscopy (980 nm) (Figure 1B−D, Supplementary Figure 1H,I). Small spines located on the first to third branches of the apical dendrites were selected for plasticity induction. Caged glutamate (4 mM CDNI-Glu) was locally applied through a pressure pipette, and a two-photon laser was directed to the tip of the spine head (720 nm). Uncaging stimulation of 3 or 4 spines, which elicited excitatory postsynaptic potential, was paired with action potentials by current injections following a spike-timing dependent plasticity (STDP) stimulation protocol (Figure 1E, Supplementary Figure 1J) (71, 72).

The STDP stimulation caused a transient increase in spine volume that returned to baseline by 50 minutes in adolescent mice (Figure 1F; n = 5 dendrites, including 19 spines from 4 mice), but it was sustained by 50 minutes in juvenile mice (2−3 weeks of age) (Figure 1G; n = 7 dendrites, including 28 spines from 3 mice). Ablation of microglia restored sustained sLTP in adolescent mice to levels similar to those in juveniles, while having little effect in the juvenile group (Figure 1H−K; clodronate in adolescent, n = 7 dendrites, including 27 spines from 5 mice; clodronate in juvenile, n = 6 dendrites, including 20 spines from 3 mice). Neighboring spines remained stable, indicating that microglial ablation preserved stimulation-specific sLTP (Figure 1H, I). A second microglia-ablating compound, PLX3397 (73), replicated the effect of clodronate in adolescent mice (Supplementary Figure 1K−M; 6 dendrites, including 23 spines from 3 mice).

Plotting spine volume changes against age revealed an age-dependent decline in sLTP in the presence of microglia (Figure 1L; w/o clodronate, data shown in Figure 1F,G, and additional data from mice aged 3−5 weeks; n = 16 dendrites, including 64 spines from 12 mice in total). In contrast, this decline was not observed when microglia were absent (w/ clodronate, data shown in Figure 1H,I; n = 13 dendrites, including 47 spines from 8 mice in total). Given the established correlation between dendritic spine volume and glutamate sensitivity in the neocortex (36, 37), these findings align with the previously reported developmental decline in LTP (14, 18–20) during the transition from juvenile to adolescent stages. Our data further suggest that microglia acquire a suppressive role during this developmental window, whereas pyramidal neurons maintain an intrinsic capacity for sLTP.

The absence of sLTP in adolescents might depend on stimulation strength. To examine this possibility, we performed plasticity induction in magnesium-free extracellular solution (Supplementary Figure 2A), which evokes a strong Ca^2+^ influx upon glutamate uncaging without requiring action potentials, a widely adopted method for sLTP induction in the hippocampus (34, 74, 75). Even under this magnesium-free condition, glutamate uncaging (0.6 ms pulse, 5 Hz, 300 times) failed to induce sLTP in slices from adolescent mice (Supplementary Figure 2B,C; 7 dendrites, including 32 spines from 5 mice). In contrast, sLTP was induced in microglia-ablated slices (Supplementary Figure 2B,D,E; 5 dendrites, including 18 spines from 4 mice), replicating the results obtained with STDP stimulation (Figure 1J).

The same magnesium-free protocol was sufficient to induce sLTP in the spines of spiny-projection neurons in the nucleus accumbens, which is located in the same coronal section as the mPFC in adolescent mice (Supplementary Figure 2F,G; 5 dendrites, including 19 spines from 4 mice). Given that the STDP stimulation protocol in the nucleus accumbens requires G_s_ activation through dopamine or adenosine for sLTP induction (71, 72), the ability of the magnesium-free condition to induce sLTP supports the view that this stimulation protocol is stronger than the STDP protocol. In contrast to the mPFC of adolescent mice, microglial ablation did not further promote sLTP in the nucleus accumbens (Figure 1M, Supplementary Figure 2F,H; 5 dendrites, including 19 spines from 4 mice), indicating regional differences in the microglial suppression of plasticity.

Acute slice preparation may alter microglial status (76). A previous study reported a gradual increase in tissue ATP levels over 5 hours after slice preparation, leading to morphological changes in microglia through receptors such as P2Y_12_ receptors (76). Because we observed microglia ablation effects only in the adolescent mPFC, it is possible that microglia in the adolescent PFC are more sensitive to ATP-related alterations than those in the nucleus accumbens or in the juvenile PFC. We therefore reanalysed the STDP stimulation results by dividing the data into the first 2.5 hours and the later period. However, we found no significant difference between these two time windows (Supplementary Figure 3A). Additionally, inhibition of P2Y_12_ receptors did not induce plasticity with STDP stimulation alone (Supplementary Figure 3B; 7 dendrites, including 27 spines from 5 mice). These observations indicate that alterations in microglia caused by acute slice preparation cannot by themselves account for the suppressive effect observed in the adolescent mPFC.

Together, these results indicate that during the transition from juvenile to adolescent stages, microglia in the mPFC acquire a suppressive function that limits Hebbian pairing induction of sLTP, whereas pyramidal neurons retain their intrinsic capacity for plasticity.

## Downstream signaling of microglial suppression

Having established that microglia suppress sLTP during adolescence, we next asked how this suppression is implemented at the cellular and molecular levels, either through direct microglial contact with spines or through the release of humoral factors (49, 54, 77–84). If suppression requires physical contact, STDP stimulation should recruit microglial processes to the vicinity of stimulated spines. In acute slices from Tmem119-CreERT2; Ai14 (Figure 2A), in which more than 99% of microglia express tdTomato (Supplementary Figure 1F), microglial protrusions were dynamically recruited to the site of adenosine triphosphate (ATP) application via a pipette within 10 minutes (Figure 2B,C), consistent with the previous report (85). When dendritic spines and microglia were imaged simultaneously, fewer than 25% of spines were located near microglial processes before stimulation (Supplementary Figure 4A−C). Unlike ATP application, STDP stimulation did not recruit the microglial processes to stimulated spines, as assessed by fluorescence intensity (Figure 2C,D) or contact counts (Supplementary Figure 4C). These results therefore do not support a mechanism involving physical contact.

**Figure 2.**
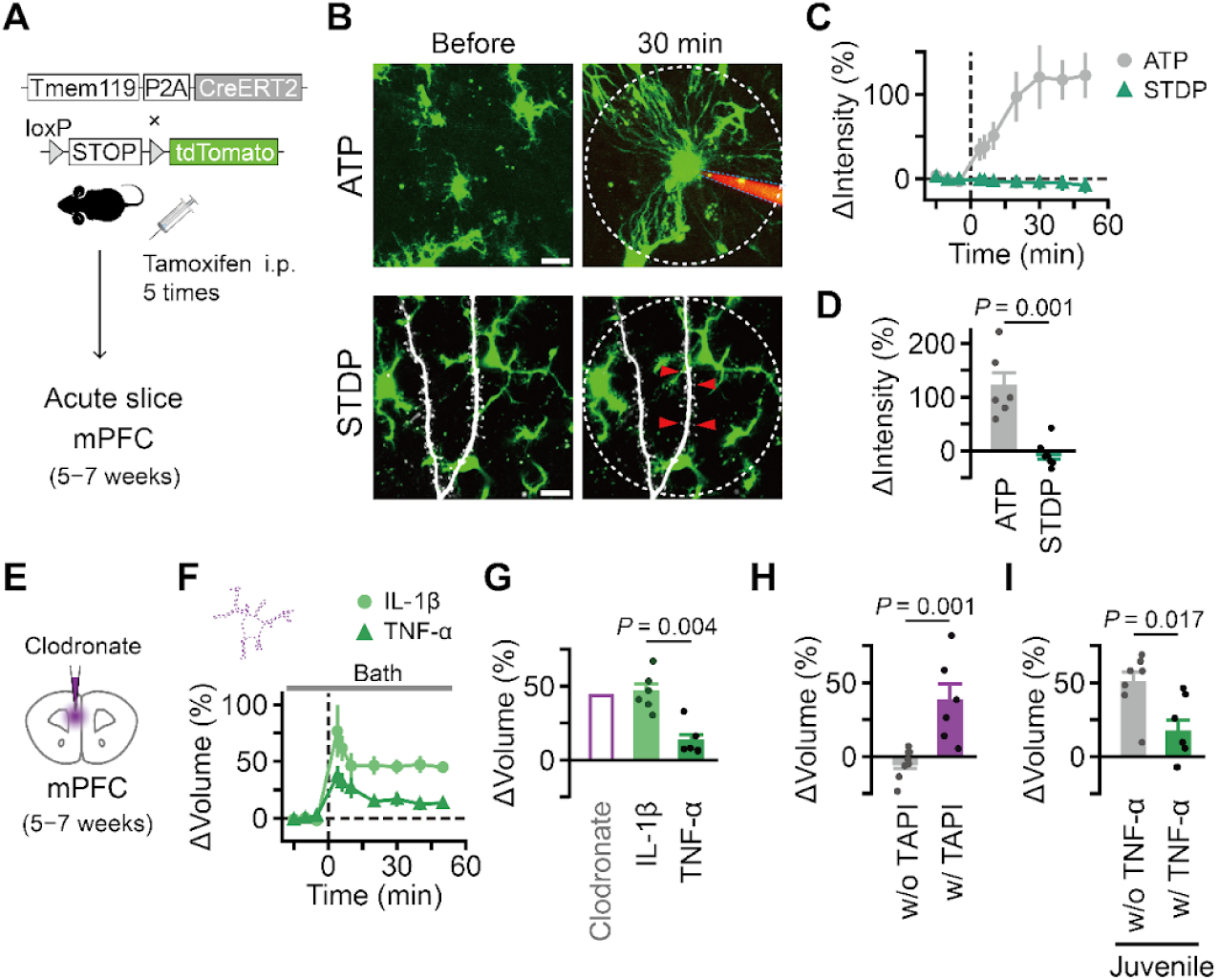
Signaling of microglial suppression. **A,** Schematic of the experimental timeline. **B,** Representative z-stack images (30 μm thick) of acute slices with 2-photon microscopy before (left) and after (right) ATP application from pipette (top) or STDP stimulation (bottom). Fluorescences of tdTomato for microglia (green), Alexa 488 for neurons (white), and Alexa 488 with 20 mM ATP in ACSF contained in the pipette (red) are shown. Red arrows indicate stimulated spines. White dotted circles denote the regions of interest (70 μm in diameter, centered at the ATP pipette tip or the stimulated spine area) for quantitative analyses. Scale bars, 10 μm. **C,D,** Time courses (C) and plots (D) of changes in intensity in the regions of interest indicated in (B). **E,** Schematic of clodronate injection. **F,G,** Time courses (F) and plots (G) of spine volume changes in the slices with clodronate injections in the presence of IL-1β and TNF-α. The clodronate injection condition shown in (Figure 1J) is indicated as a reference. **H,I,** Plots of spine volume changes following STDP stimulation in the absence and presence of TAPI-0 (TACE/ADAM17 inhibitor) (H) or in the absence and presence of TNF-α from juvenile mice (I). No clodronate injections were performed in these experiments. Error bars represent S.E.M. Mann–Whitney U test used for statistical analysis in (D,G,H,I).

We thus tested whether humoral factors—tumor necrosis factor-α (TNF-α) and interleukin-1β (IL-1β) —previously suggested to suppress LTP in the hippocampus (80–82) and preferentially expressed in the microglia in the neocortex (86–90), would also suppress sLTP in the mPFC. In microglia-ablated slices (Figure 2E), bath applied TNF-α (10 nM, bath) (91), but not IL-1β (1 ng/ml, bath) (81), suppressed STDP-induced sLTP (Figure 2E−G, Supplementary Figure 4D−F; TNF-α, n = 6 dendrites, including 21 spines from 4 mice; IL-1β, n = 6 dendrites, including 21 spines from 4 mice). TNF-α can signal through a soluble form released after cleavage by TNF-α converting enzymes (TACE) or A disintegrin and metalloprotease 17 (ADAM17) (92, 93). Inhibition of TACE/ADAM17 by TAPI-0 (1 µM, bath) allowed sLTP in the presence of microglia (Figure 2H, Supplementary Figure 5G; w/o TAPI-0, n = 8 dendrites, including 31 spines from 4 mice; w/ TAPI-0, n = 6 dendrites, including 17 spines from 5 mice). Another class of TNF-α inhibitor, pomalidomide (100 µM, bath) (94, 95), further supported the involvement of TNF-α signaling (Supplementary Figure 5H; n = 6 dendrites, including 21 spines from 4 mice). Moreover, TNF-α suppressed sLTP even in the juvenile mPFC (Figure 2I, Supplementary Figure 4I; n = 7 dendrites, including 26 spines from 5 mice).

Given that microglia are a major source of TNF-α (88–90), these results suggest that microglial suppression of sLTP involves the humoral action of TNF-α in adolescent mice, potentially reflecting developmental changes in microglial properties (96).

## Microglia-mediated actions of NA on spine enlargement

We next tested whether neuromodulatory signals could relieve the microglial suppression under physiological conditions, thereby enabling a three-factor rule for sLTP induction. NA was a strong candidate, given its established role in modulating neocortical synaptic plasticity (13, 27, 97) and its direct effects on microglial function (51, 52). We therefore locally delivered NA (50 µM) via a pressure pipette along with the caged-glutamate, beginning 3−5 minutes before the onset of STDP stimulation to mimic the transient NA release observed *in vivo* (62) (Figure 3A−G). The presence of NA during STDP stimulation enhanced sustained sLTP in the stimulated spines up to 50 minutes (Figure 3B,C,G; STDP only, n = 9 dendrites, including 36 spines from 6 mice; STDP with NA, n = 7 dendrites, including 22 spines from 6 mice; *P =* 0.028, post hoc Steel’s test against STDP after Kruskal–Wallis test against the following experiments shown in Figure 3C−G, Supplementary Figure 5A−H). In contrast, dopamine (50 µM, puff), another catecholamine known to influence spine dynamics *in vivo* (98), did not promote sLTP, unlike its effect in the nucleus accumbens (71) (Figure 3G, Supplementary Figure 5A; n = 6 dendrites, including 24 spines from 4 mice; *P =* 1.00).

**Figure 3.**
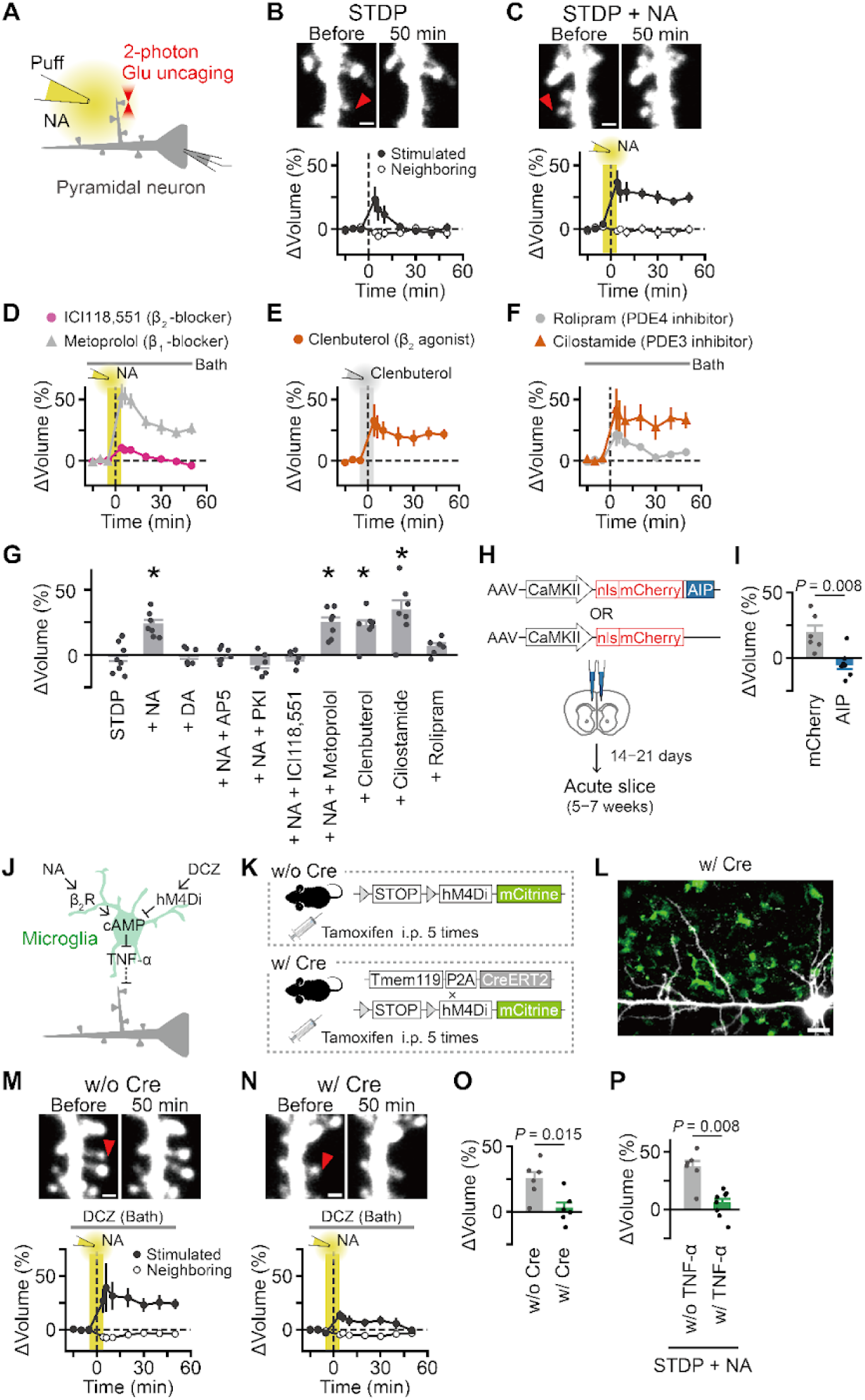
Microglia-mediated noradrenaline actions on spine enlargement. **A,** Schematic of the experimental configuration for puff application and two-photon uncaging on dendritic spines of a pyramidal neuron. **B,C,** Representative images of dendritic spines before and 50 minutes after STDP stimulation (top) and time courses of spine volume changes after STDP stimulation (bottom) in the absence (B) and presence (C) of NA. Red arrows indicate stimulated spines. Shaded areas indicate the period of puff application. Scale bars, 1 μm. **D–F,** Time courses of spine volume changes in the presence of β blockers with NA (D), β_2_ agonist (E), and phosphodiesterase inhibitors (F). Gray shade and bars indicate the period of drug application. **G,** Plots of spine volume changes. Volume changes were averaged over 40−50 minutes and subjected to statistical tests (Kruskal–Wallis test and post hoc Steel’s test against STDP only). \**P* < 0.05. Error bars represent S.E.M. **H,** Schematics of AAV vectors to express mCherry with or without AIP. **I,** Plots of spine volume changes. **J,** Schematic of a signaling model. **K,** Schematic of R26-LSL-hM4Di-mCitrine mice without (w/o Cre) and with (w/ Cre) Tmem119-CreERT2 allele. **L,** Image of an acute slice with 2-photon microscopy from a transgenic mouse (w/ Cre). Fluorescences of mCitrine (green) and Alexa 594 (white) are shown. Scale bar, 20 µm. **M,N,** Representative images of dendritic spines before and 50 minutes after STDP stimulation (top) and time courses of spine volumes (bottom) in the slices without (M) and with (N) Cre. Gray bars indicate the period of DCZ (100 nM) application. Red arrows indicate stimulated spines. Scale bars, 1 μm. **O,** Plots for spine volumes shown in (M,N). **P,** Plots of spine volume changes in the absence and presence of TNF-α following STDP stimulation in the presence of NA from Supplementary Figure 6G. Error bars represent S.E.M. Statistical analysis was performed using the Mann–Whitney U test in (F,G).

Next, we characterized downstream signaling of NA by pharmacologically perturbing G_s_-coupled β adrenergic receptors (ARs), which stimulate cAMP production, and phosphodiesterase (PDE), which degrades cAMP, given that the β-ARs-cAMP-PKA pathway generally facilitates LTP (99, 100). RNA-seq data provide comprehensive information on the cellular distribution of β-ARs in the neocortex (86, 101), showing that β_1_- and β_3_-ARs are primarily localized to neurons and astrocytes, whereas β_2_-ARs are predominantly expressed in microglia. Bath application of ICI118,551 (1 μM, β_2_ inverse agonist) blocked NA-enhanced sLTP (Figure 3D,G, Supplementary Figure 5B, n = 6 dendrites, including 23 spines from 3 mice; *P =* 1.00), whereas metoprolol (1 μM, β_1_ inverse agonist) did not (Figure 3D,G, Supplementary Figure 5C, n = 7 dendrites including 28 spines from 5 mice; *P =* 0.038). Moreover, puff application of clenbuterol (10 μM, a β_2_ agonist) reproduced the effect of NA on sLTP (Figure 3E, Supplementary Figure 5D, n = 7 dendrites, including 23 spines from 7 mice; *P =* 0.028). Among the PDE families, PDE3b is preferentially expressed in microglia (102), whereas PDE4 subtypes are expressed in pyramidal neurons (86). We therefore hypothesized that PDE inhibition could substitute for G_s_-coupled receptor stimulation. STDP alone induced sLTP in the presence of cilostamide (1 μM bath, PDE3 inhibitor) (Figure 3F,G, Supplementary Figure 5E; n = 7 dendrites, including 27 spines from 5 mice; *P =* 0.028). In contrast, rolipram (1 μM bath, PDE4 inhibitor), at a concentration known to enhance hippocampal LTP (103), did not facilitate sLTP (Figure 3F,G, Supplementary Figure 5F; n = 6 dendrites, including 24 spines from 3 mice; *P =* 0.84). These pharmacological results support the hypothesis that NA-mediated sLTP facilitation involves β_2_-adrenergic receptors on microglia.

Further pharmacological experiments confirmed that the NA-dependent sLTP was abolished by D-AP5 (50 µM, bath application; NMDA receptor antagonist) (Figure 3G, Supplementary Figure 5G; n = 7 dendrites, including 27 spines from 6 mice; *P =* 0.99) and by PKI (10 µM, a PKA inhibitor, in the intracellular solution) (Figure 3G, Supplementary Figure 5H; n = 6 dendrites, including 23 spines from 4 mice; *P =* 0.98). We next tested whether expressing the CaMKII inhibitory peptide AIP (autocamtide-2-related inhibitory peptide) prevents sLTP (71, 104). AIP was delivered using an AAV vector under CaMKII promoter (Figure 3H). NA-dependent sLTP was observed in neurons expressing the control vector (Figure 3I, Supplementary Figure 5I; n = 6 dendrites, including 24 spines from 6 mice) but was blocked in AIP-expressing neurons (Figure 3I, Supplementary Figure 5J, n = 7 dendrites, including 23 spines from 4 mice). These data indicate that the essential signaling mechanisms underlying sLTP are similar to those described in the hippocampus and striatum (42, 71, 72).

Using microglia-specific chemogenetic inhibition of cAMP signaling, we further tested whether microglia mediate the action of NA (Figure 3J). We used double-transgenic mice expressing hM4Di, a G_i/o_-coupled receptor that suppresses cAMP production in a Cre-dependent manner (R26-LSL-hM4Di-mCitrine)(105) under the control of Tmem119-CreERT2 (Figure 3K). This combination has been used in recent studies to modulate microglial functions (106, 107). The R26-LSL-hM4Di-mCitrine line is known to exhibit leak expression, which was observed in 3% of cells in the mPFC (Supplementary Figure 6A−C). After repeated intraperitoneal tamoxifen injections (Supplementary Figure 6D), we confirmed expression of the chemogenetic probe in microglia in fixed slices, along with leak expression in other cell types (Supplementary Figure 6E,F).

In acute slices, we recorded from pyramidal neurons without detectable mCitrine fluorescence and applied deschloroclozapine (DCZ, 100 nM), a potent hM4Di actuator(108–110) in the bath. We found that the presence of Cre prevented sLTP induction by STDP stimulation with NA (Figure 3L−O; without Cre, n = 6 dendrites, including 24 spines from 3 mice; with Cre, n = 6 dendrites, including 24 spines from 4 mice). Importantly, STDP-induced sLTP was fully preserved in non-Cre slices from the same double-transgenic mice (Figure 3C vs. Figure 3M), indicating that leak expression and potential off-target effects of DCZ did not impair plasticity. These results therefore indicate that microglial cAMP signaling mediates the NA-dependent enhancement of spine enlargement.

Microglial ablation experiments suggested that TNF-α suppresses plasticity downstream of microglia (Figure 2G, Figure 3J), implying that TNF-α may interfere with the effect of NA on plasticity. Consistent with this idea, TNF-α application reduced NA-enabled sLTP induced by STDP (Figure 3P, Supplementary Figure 6G, w/o TNF-α, n = 6 dendrites, 24 spines from 5 mice; w/ TNF-α, n = 8 dendrites, 31 spines from 5 mice).

Collectively, our *ex vivo* experiments demonstrate that microglia play a key role in the developmental transition of plasticity rules from a Hebbian form to a three-factor form for sLTP induction in the mPFC. In this framework, microglia adopt a suppressive role in Hebbian plasticity after maturation through a humoral factor, whereas NA can counteract this suppression via β_2_-ARs and cAMP signaling within microglia.

## NA-dependent plasticity signaling in rapid social learning

We next tested whether the three-factor rule governed by the NA-dependent plasticity pathway contributes to social emotional learning using the observational fear learning (OFL) paradigm, which engages the mPFC (64, 65, 69, 111). Pairs of male adolescent mice (6–7 weeks old) were co-housed for 4 days. On day 1, one mouse (the observer) received two foot shocks (0.4 mA, 2 s) in a conditioning chamber (Figure 4A). On day 2, the cage mate (the demonstrator) was placed in the same chamber and exposed to 24 tone–shock pairings (6 kHz tone, 5 s; 0.7 mA shock, 2 s). Tone and shock presentations elicited transient freezing in observers regardless of demonstrator presence, likely due to novelty, whereas sustained freezing occurred only when observers witnessed their cage mate being shocked (Figure 4B, Supplementary Figure 7A; with demonstrator, n = 8; without, n = 6). On day 3, memory was assessed by presenting the tone alone (6 kHz, 60 s, 3 trials). Observer mice exhibited robust freezing during tone presentation, but only if they had previously witnessed the demonstrator being shocked (Figure 4C, Supplementary Figure 7B). Although female mice also acquired OFL, their freezing responses on day 2 were weaker (Supplementary Figure 7C–F, n = 6). Because day 2 freezing was used as an internal reference for subsequent experiments, females were excluded from further testing.

**Figure 4.**
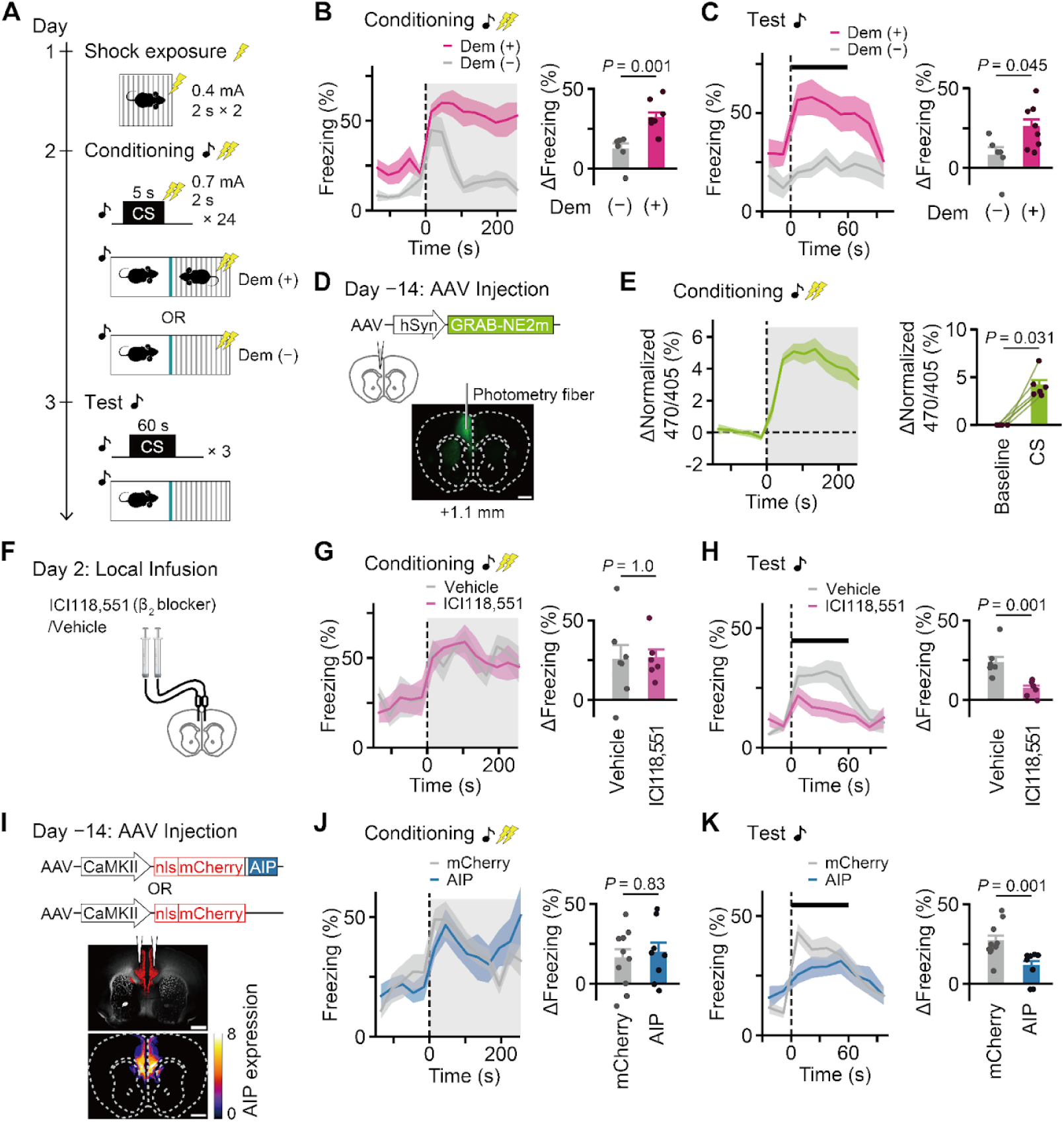
Noradrenaline and plasticity-related signaling in observational fear learning. **A,** Schematic of the experimental timeline of observational fear learning. **B,** Traces (left) and plots (right) of freezing on day 2. Grey shade indicates the conditioning period. The changes in freezing during the conditioning periods compared to the baseline periods on day 2 were plotted. **C,** Traces (left) and plots (right) of freezing on day 3. Averaged freezing responses over the CS presentations are shown, and the bar indicates the CS period. The changes in freezing during the CS periods compared to the baseline periods on day 3 were plotted. **D,** Schematic of the virus injection and fiber photometry experiment in the mPFC (top) and representative image showing fluorescence of GRAB-NE_2m_ (green) (bottom). The trace of fiber was overlaid on the image. Scale bar, 1 mm. **E,** Traces (left) and plots (right) of GRAB-NE_2m_ fluorescence on day 2. **F,** Schematic of cannula injection of ICI118,551 (β_2_ inverse agonist). The experimental timeline of observational fear learning is the same as (A). **G, H,** Traces and plots of freezing on day 2 (G) and on day 3 (H) in mice with injections of ACSF or ICI118,551 on day 2. **I,** Images showing the spread of AIP expression. Representative (top) and the average of mCherry fluorescences after binarization from all the mice used in behavioral experiments (n = 8 mice) are shown. Scale bar, 1 mm. **J, K,** Traces and plots of freezing on day 2 (J) and on day 3 (K) in mice injected with mCherry or AIP. Mann–Whitney U test for (B, C, G, H, J, K). Wilcoxon signed-rank test for (E). Error bars and shades indicate S.E.M.

We then assessed NA dynamics in the mPFC using the GPCR-based fluorescent sensor GRAB-NE_2m_ (112). An AAV expressing GRAB-NE_2m_ under the synapsin promoter was injected into the right mPFC, and fluorescence was recorded through an implanted optical fiber (Figure 4D). As a positive control, direct foot shock elicited a transient fluorescence increase that peaked within seconds and decayed over tens of seconds (Supplementary Figure 7G,H; *n* = 5 mice), consistent with prior reports (61). During OFL, GRAB-NE_2m_ signals in observer mice increased transiently during the 240-second shock period delivered to demonstrator mice on day 2 (Figure 4E, Supplementary Figure 7I, n = 6 mice), consistent with noradrenergic axon activity patterns (62). In contrast, no signal increase was observed on day 3 during cue-driven recall (Supplementary Figure 7J–L). These results indicate that demonstrator distress evokes a sustained, minute-scale NA response in the mPFC, resembling the timing of NA application in our slice experiments (Figure 3A,C). Although GRAB-NE_2m_ was expressed in neurons, this volume-transmitted NA signal likely reflects levels accessible to microglia.

To test the role of β_2_-ARs in the acquisition of OFL, we bilaterally infused ICI118,551 or ACSF as a vehicle control into the mPFC on day 2 through pre-implanted cannulas (Figure 4F). Infusion of ICI118,551 did not alter freezing response during conditioning but subsequently inhibited freezing on day 3 (Figure 4G,H, Supplementary Figure 7M,N, Vehicle, n = 7 mice; ICI118,551, n = 6 mice). Because freezing during conditioning typically correlates with memory strength (67), this dissociation suggests that β_2_-AR blockade selectively impaired memory formation rather than fear expression. Together with reports that non-selective β-AR antagonists suppress vicarious fear responses (62), these findings indicate that β_2_-AR in the mPFC preferentially contributes to memory formation, whereas β_1_-AR may play a more direct role in fear expression.

Given that AIP prevented sLTP induction, we bilaterally expressed AIP in the mPFC and anterior cingulate cortex (ACC) (Figure 4I, Supplementary Figure 8A). This manipulation did not alter freezing during conditioning on day 2 (Figure 4J, Supplementary Figure 8B, mCherry, n = 10 mice; AIP, n = 8 mice) but impaired freezing on day 3 test (Figure 4K, Supplementary Figure 8C), consistent with the effects observed in the β_2_ blockade experiment.

These results indicate that the NA-dependent plasticity pathway in the mPFC is required for the rapid formation of social fear learning in adolescent male mice, supporting a role for the three-factor rule in plasticity induction during emotional learning.

## Microglial role in rapid social learning

As microglia suppress sLTP induction in the adolescent mPFC (Figure 1J), we predicted that microglial ablation in the mPFC would enhance the acquisition of observational fear learning (OFL). Clodronate was unilaterally injected into the right mPFC on day 0 through a glass pipette, and its spread extended into the anterior ACC by the following day (Figure 5A–C, Supplementary Figure 9A). To avoid ceiling effects during OFL, demonstrator mice received lower-intensity foot shocks (0.2 mA) with fewer repetitions (12 trials) on day 2 (Figure 5A). Under this subthreshold protocol, control observer mice showed freezing during conditioning (Figure 5D, Supplementary Figure 9B) but showed only modest freezing to the tone during testing on day 3 (Figure 5E, Supplementary Figure 9C). Microglial ablation did not affect freezing during conditioning (Figure 5D, Supplementary Figure 9B; Vehicle, *n* = 7 mice; Clodronate, *n* = 6 mice) but significantly increased freezing on day 3 (Figure 5E, Supplementary Figure 9C). These results support the interpretation that microglial ablation facilitates learning rather than simply enhancing threat sensitivity.

**Figure 5.**
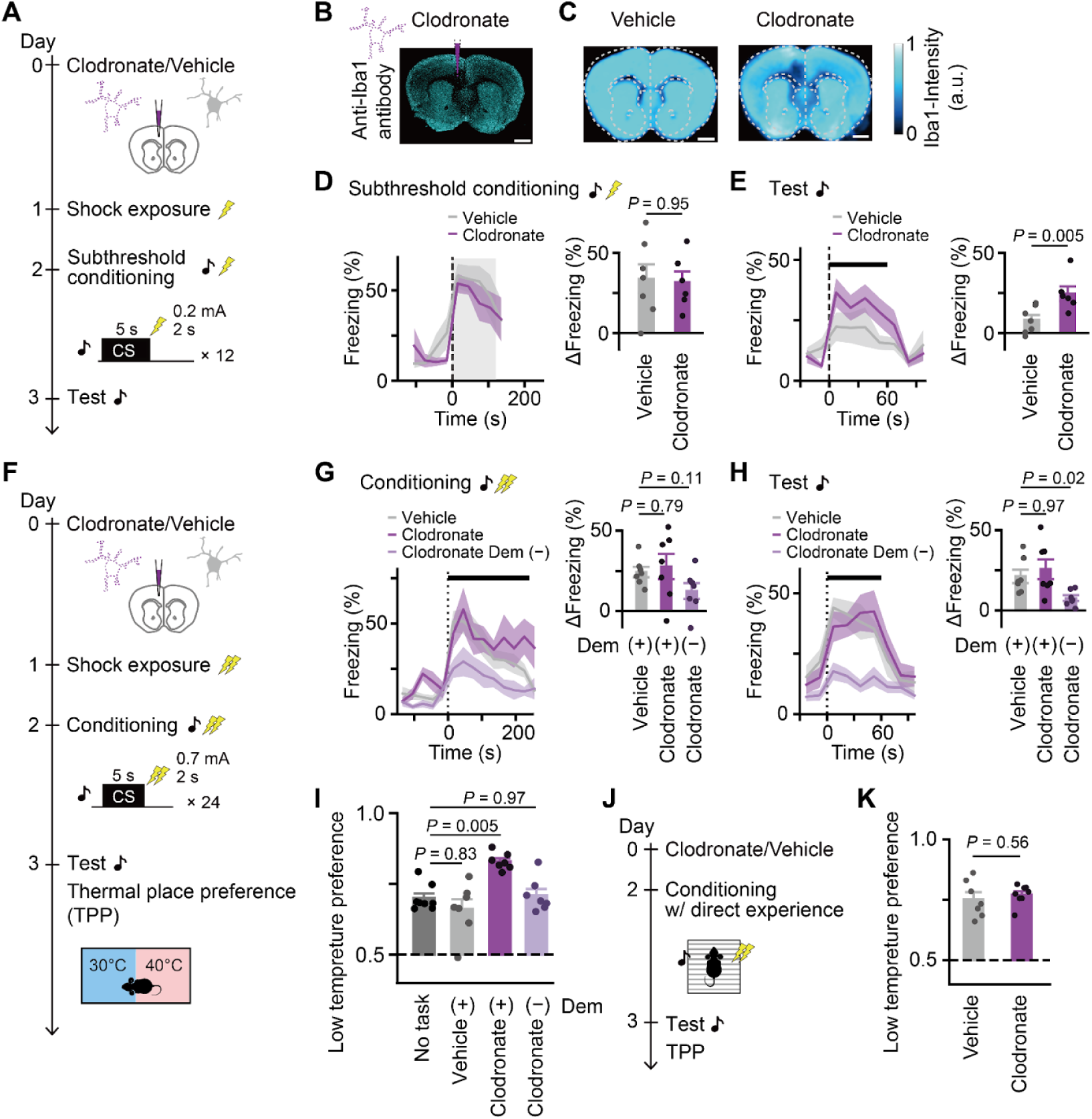
Microglial protection of cross-talk between emotional circuits. **A,** Schematics of experimental timeline for subthreshold observational fear learning. **B,** Representative image showing Iba1 immunoreactivity in clodronate injected brain. Scale bars, 1 mm. **C,** Averaged images showing Iba1 immunoreactivity in the vehicle (left) and clodronate (right) injected brain. Averaged images were generated from all the brains used in behavioral experiments shown in (D) and (E). Dark color indicates areas with low Iba1 immunoreactivity. Scale bars, 1 mm. **D, E,** Traces and plots of freezing on day 2 (D) and on day 3 (E) in mice with injections of vehicle or clodronate in subthreshold observational fear learning. **F,** Schematic of experimental timeline for observational fear learning and subsequent thermal place preference test. **G, H,** Traces and plots of freezing on day 2 (G) and on day 3 (H) in mice with injections of vehicle or clodronate in observational fear learning, and in mice with injections of clodronate without demonstrator. **I,** Plots of thermal preference place after observational fear learning. Data for “No task” are the same as that shown in Supplementary Figure 9D. **J,** Schematic experimental timeline for direct fear learning and subsequent thermal place preference test. **K,** Plots of thermal place preference test after direct fear learning. Error bars and shades indicate S.E.M. Mann–Whitney U test for (D, E, K), Kruskal–Wallis test and post hoc Steel’s test for (G, H, I). *P* = 0.13, 0.0027, and 0.0027 for Kruskal–Wallis test in (G, H, I).

To examine the functional role of microglial suppression of sLTP, we next asked whether enhanced plasticity in the absence of microglia leads to the recruitment of task-irrelevant, emotion-related circuits. Pain is an emotional function partly regulated by the mPFC, including the ACC, where synaptic potentiation modulates sensitivity (113, 114). Notably, some neurons in this region encode overlapping representations of pain and vicarious fear (68), suggesting that precise plasticity control is required to update vicarious fear memories without disrupting pain-related processing. We therefore hypothesized that OFL in the absence of microglia could alter pain sensitivity in a learning-dependent manner. To assess pain sensitivity, we used the thermal place preference (TPP) test (66), in which mice were allowed to freely explore two plates maintained at different temperatures. Avoidance of the warmer plate serves as a readout of thermal nociception. Control mice showed a modest tendency to avoid 40 °C plate, no discrimination between identical 30 °C plates, and consistent avoidance of the 50 °C plate (Supplementary Figure 9D; *n* = 8 mice).

We repeated OFL using the standard protocol to compare animals with and without microglial ablation in the right mPFC (Figure 5F). Freezing levels on both day 2 and day 3 were comparable between groups, consistent with a ceiling effect under these conditions (Figure 5G, H, Supplementary Figure 9E–G; Vehicle, *n* = 7 mice; Clodronate, *n* = 7 mice). We then assessed thermal sensitivity using the TPP test (30 °C vs. 40 °C) and found that vehicle-injected mice did not differ from naïve controls (“No task”), indicating that OFL alone did not alter thermal nociception (Figure 5I). In contrast, mice that underwent OFL in the absence of microglia (clodronate-injected) showed significantly increased thermal aversion (Figure 5I). This effect was absent in clodronate-injected mice exposed to the task chamber without a demonstrator (Figure 5G–I, Supplementary Figure 9F, G; Clodronate Dem (−), *n* = 7 mice). To confirm the involvement of plasticity-related signaling in thermal sensitivity, we unilaterally inhibited CaMKII signaling by injecting AIP and subsequently ablated microglia on the same side (Supplementary Figure 10A–C). AIP expression occluded the clodronate-induced enhancement of thermal sensitivity after OFL under the standard protocol (Supplementary Figure 10D–J; AIP and Vehicle, *n* = 6 mice; AIP and Clodronate, *n* = 6 mice). These results support that the microglial ablation enhances thermal sensitivity through plasticity-dependent mechanisms. Together, these findings demonstrate that, in the absence of microglia, social fear learning can aberrantly amplify pain sensitivity, leading to experience-dependent cross-talk between emotional circuits and highlighting a functional role for mature microglia in stabilizing established circuits during learning.

In contrast to the rapid remodeling of the mPFC network during social learning (69), fear learning based on direct experience induces slower remodeling of the mPFC (115), suggesting a lower demand on rapid plasticity. To confirm this, we performed auditory fear conditioning using direct fear experience. Mice received four tone–shock pairings on day 1 (Supplementary Figure 8D), followed by tone presentations on day 2. Unlike its effect on OFL, CaMKII inhibition through AIP expression in the mPFC did not impair the rapid acquisition of fear conditioning from direct experience (Supplementary Figure 8E, F; mCherry, *n* = 6 mice; AIP, *n* = 5 mice), consistent with previous evidence of delayed synaptic remodeling in the mPFC (115). Similarly, microglial ablation in the right mPFC did not affect conditioning or subsequent thermal sensitivity (Figure 5J, K, Supplementary Figure 9H, I; Vehicle, *n* = 7 mice; Clodronate, *n* = 8 mice). These findings highlight a selective requirement for finely tuned plasticity mechanisms to support rapid social learning without inducing emotional cross-talk.

## Discussion

We found that microglia play a key role in the developmental shift of plasticity rules from a Hebbian plasticity form to a three-factor plasticity form in the mPFC during the transition from juvenile to adolescent stages. After maturation, microglia acquire the ability to suppress Hebbian plasticity through a humoral factor, and this suppression is counteracted by β_2_-adrenergic receptor–mediated cAMP signaling, enabling sLTP and rapid social learning. Based on these findings, we propose a microglia-mediated three-factor rule in which microglia gate synaptic potentiation by requiring noradrenergic input as a third factor. Moreover, this microglia-mediated suppression of Hebbian plasticity prevents social fear learning from spilling over into heightened pain sensitivity, suggesting that microglia tune the balance between plasticity and rigidity to preserve emotional circuit specificity in the mature mPFC.

We isolated the postsynaptic component of a neocortical Hebbian plasticity rule at single spine level using two-photon glutamate uncaging during the transition from juvenile to adolescent stages. We confirmed its developmental decline of sLTP across this period, consistent with previous reports (14, 19–22). In contrast to earlier models attributing this decline to neuron-intrinsic mechanisms (18–20, 116, 117), our results show that microglia play a critical role, likely through the humoral release of TNF-α, while pyramidal neurons retain their plastic potential at least until 7 weeks of age (Figure 1H). Although we cannot fully exclude the possibility that microglial suppression of sLTP reflects artefacts related to slice preparation, behavioral experiments involving β_2_-AR inhibition and microglial ablation support the microglial suppression model. Unlike inhibitory mechanisms that irreversibly constrain plasticity after the critical period closure (118), microglial suppression is unique in that it can be rapidly reversed by a single physiological signal of NA.

In the adolescent mPFC, a three-factor rule for sLTP induction is achieved through NA, which counteracts microglia-dependent suppression, extending prior findings on the role of NA/β-AR/cAMP/PKA signaling in promoting neocortical LTP (13, 27, 97, 119–121). Consistent with earlier studies showing that neuronal cAMP/PKA signaling is essential for LTP (122) and sLTP (71, 123), we found that NA-dependent neocortical sLTP required PKA activity in pyramidal neurons, as evidenced by the blockade of sLTP when the PKA inhibitor PKI was included in the recording pipette (Figure 3G). Since neuronal PKA is canonically driven by synaptic activity and calcium influx (124), these results support a dual-regulation model: while neuronal PKA drives the intracellular signaling for sLTP induction, this machinery is constitutively clamped by a microglial brake; Consequently, neuronal PKA activation alone was insufficient to induce sLTP in the mature mPFC; β_2_-ARs/cAMP signaling in microglia was additionally required to relieve this suppression. In contrast to the facilitatory role of microglial β_2_-ARs in governing the three-factor rule for sLTP induction, a previous *in vivo* study in the visual cortex reported a suppressive role for microglial β_2_-ARs in spine remodeling through regulation of microglial surveillance over the course of days (52). These findings suggest that microglial β_2_-ARs may operate on two distinct timescales to control spine remodeling in opposing directions.

A critical question concerns the timescale of NA and TNF-α dependent modulation. Given that TNF-α mediates plasticity suppression, NA may act by reducing TNF-α release, consistent with its known immunomodulatory effects in monocytes (125, 126) and macrophages (127, 128). Alternatively, NA may trigger the release of a neutralizing factor that counteracts TNF-α. In either case, it is crucial whether the suppression of TNF-α can explain the rapid recovery of sLTP observed within minutes of NA application. Although real-time monitoring of endogenous TNF-α or downstream dynamics at the synapse level remains technically challenging, several lines of evidence support the physiological plausibility of this rapid switching. First, GPCR-mediated cAMP signaling is generally immediate, occurring within seconds (129). Second, the synaptic actions of TNF-α are known to be highly dynamic; previous studies have established that TNF-α can regulate AMPA receptor trafficking and synaptic efficacy within 10 minutes (91, 130). This suggests that the brake imposed by TNF-α is not a static structural constraint but a continuously active signaling pressure that, once removed, allows the synaptic machinery to engage quickly. Thus, despite the methodological difficulty in directly visualizing the rapid dynamics of extracellular TNF-α, the rapid relief of microglial suppression provides a plausible explanation for the gating of plasticity rules.

The NA, β_2_-ARs, and CaMKII-dependent signaling for the three-factor rule appears to contribute to the rapid establishment of social learning in the mPFC. Our data supports that the mPFC undergoes active synaptic remodeling during observational learning (69), rather than merely relaying social information. In contrast, this pathway was dispensable for learning from direct experience, consistent with prior reports showing that mPFC synaptic potentiation emerges only days after direct conditioning (115). NA levels rise in the mPFC during both observation and direct shock (Figure 4E, Supplementary Figure 7H), indicating that NA release alone does not account for the selective engagement of plasticity during social learning. Instead, we propose that distinct patterns of glutamatergic activity during social observation uniquely recruit mPFC plasticity. Supporting this idea, microglial ablation enhanced pain sensitivity only after social, but not direct, fear learning (Figure 5I, K). Crucially, this phenotype appeared without altering baseline pain thresholds or freezing behavior during acquisition, ruling out systemic hypersensitivity. Furthermore, this behavior was plasticity-dependent as inhibiting CaMKII with AIP during learning occluded the effect of microglial ablation (Supplementary Figure 10). This supports that the aberrant behavior involves unconstrained synaptic plasticity, rather than the loss of other microglial functions such as homeostatic support per se. Given that mPFC neurons often exhibit mixed selectivity (131), including overlapping representations of vicarious fear and pain-related signals (68), plasticity must be tightly constrained, as uncontrolled synaptic changes can destabilize mature networks (30, 31). By ablating microglia, we directly demonstrated that microglial gating restricts experience-driven modifications to prevent aberrant circuit cross-talk in the mature mPFC. Thus, the three-factor rule mediated by NA–microglia interactions may gate synaptic modifications within an optimal range, contributing to the functional integrity of emotional circuits.

A similar NA–microglia-dependent plasticity rule shifting may operate in other neocortical regions. Rapid spine remodeling has been observed in motor and sensory cortices *in vivo* (*38–40*), where sLTP is limited by glutamatergic stimulation alone in the visual cortex (44, 132), but not in the motor cortex (133), suggesting plasticity rule shifting in select neocortical regions including the mPFC and visual cortex. After maturation, three-factor rules may operate on different timescales across neocortical regions, as NA signaling during social fear learning is sustained over minutes in the mPFC (Figure 4E), whereas visual cortical synapses respond to NA within seconds to modulate plasticity (29, 119). Future study may clarify how the NA–microglia-dependent mechanism contributes to neocortical plasticity algorithms across regions and developmental stages. Notably, whereas microglia suppressed plasticity and learning in the mPFC, microglia in the hippocampus facilitate fear conditioning driven by direct experience (55, 134), underscoring region-specific roles of microglia in plasticity and learning rules.

In conclusion, microglia play a central role in regulating developmental transitions in synaptic plasticity rules within the mPFC. Incorporating such developmental shifts in learning rules into computational models of learning may provide new insights into how neural circuits balance plasticity and rigidity (31). Future studies will reveal molecular mechanisms in microglia controlling this transition. The transition from juvenile to adolescent stages represents a critical window for the emergence of psychiatric disorders (135), during which microglia have been increasingly implicated in disease pathophysiology (136). Elucidating roles of microglia-dependent developmental shifts in plasticity rules in pathological contexts may therefore provide a cellular framework for understanding psychiatric vulnerability.

## Acknowledgement

We thank Y. Li for the GRAB sensor plasmid; K. Hagihara, R. Hira, H. Kasai, A. Kumar, T. Masuda, J. Nagai, T. Okuyama, K. F. Tanaka, and A. Uematsu for helpful discussions; and M. Asaumi, A. Abe, A. Kurabayashi, R. Miyashita, and A. Nishikawa for technical assistance. This work was supported by Brain/MINDS (19dm0207069 to S.Y.) and Multidisciplinary Frontier Brain and Neuroscience Discoveries (Brain/MINDS 2.0) (JP23wm0625001, JP24wm0625308, JP24wm0625101 to S.Y.) from AMED; Grants-in-Aid (JP21H05171, JP21H05176, and 24K02115 to S.Y.) from JSPS; Moonshot R&D (JPMJMS2021 to S.Y.) from JST; The Uehara Memorial Foundation (to S.Y.); and The Mitsubishi Foundation (to S.Y.).

## Author contribution

S.Y. conceptualized the study. M.T. performed and A.I. and Y.I. assisted slice experiments. H.O. performed and T.S. assisted behavioral experiments. H.E. produced AAV vectors. All the authors interpreted the data. S.Y., M.T and H.O. drafted and all the authors edited the manuscript.

## Declaration of interests

The authors declare no competing interests.

## Data availability

All data needed to evaluate the conclusions in the paper are present in the paper.

## Methods

### Animals

C57BL/6J mice were purchased from Sankyo lab service. Tmem119-CreERT2 mice [C57BL/6-Tmem119 (Cre/ERT2) Gfng/J; stock #031820] (70), Ai14 mice [B6.Cg-Gt (ROSA) 26Sortm14 (CAG-tdTomato) Hze/J; stock #007914] and R26-LSL-hM4Di-mCitrine mice [B6.129-Gt (ROSA) 26Sortm1 (CAG-CHRM4*,-mCitrine) Ute/J; stock #026219] (105) were obtained from the Jackson Laboratory. Mice were housed on a 12-hour/12-hour light/dark cycle with *ad libitum* access to food and water. All animal experiments were approved by the Animal Experimental Committee of the Faculty of Medicine at the University of Tokyo.

### Plasmids and adeno-associated viruses (AAV)

For AAV production, we prepared the following constructs using PCR and Infusion (Takara): pAAV-CaMKII(0.3)-FAS-NLS-mCherry-W and pAAV-CaMKII(0.3)-FAS-NLS-mCherry-P2A-AIP-W, where FAS is loxP-FAS that has no effect in this study, NLS is nuclear localisation sequence, AIP autocamtide 2-related inhibitory peptide (KKALRRQEAVDAL), and P2A a self-cleaving peptide or obtained a plasmid from Addgene (pAAV-hSyn-GRAB-NE2m-W, #208686) (112). AAVs were produced and their titres were measured as described previously (137). Briefly, AAV expression vector, pHelper (Stratagene) and either pRepCap AAV5 (Applied Viromics) or pUCmini-iCAP-PHP.eB (#103005, Addgene) were transfected into HEK293 cells (AAV293, Stratagene). After three days, cells were harvested and AAV were purified twice using ultracentrifuge with iodixanol. The titres for AAV (genome copies, GC) were estimated using quantitative PCR.

### Surgery

For AAV injection, mice were anesthetized with isoflurane (1-3%) and placed in a stereotaxic frame (Narishige). AAVs were bilaterally (AIP experiments) or unilaterally (photometry experiments) infused into the mPFC (AP +1.8 mm, ML ±0.3 mm, DV +2.4 mm for slice experiments; AP +1.1 mm, ML ±0.3 mm, DV +2.0 mm for behavioral experiments) and using a glass needle at a rate of 100 nl/min. Either of following AAVs was infused 1 μl per hemisphere:

- AAV2/5: CaMKII(0.3)-FAS-NLS-mCherry-W (2.0 × 10^13^ GC/ml)
- AAV2/5: CaMKII(0.3)-FAS-NLS-mCherry-P2A-AIP-W (2.0 × 10^13^ GC/ml)
- AAV2/PHP.eB: hSyn-GRAB-NE_2m_-W (2.0 × 10^13^ GC/ml)

The glass needle was withdrawn 5 minutes after the end of the infusion. For photometry experiments, the optical cannula (NA 0.5, 200 μm in diameter, 3.0 mm in length, RWD) was placed in the same coordinate to the injection. The mice were allowed to recover in their home cages for at least 2 weeks before slice or behavioral experiments.

For slice experiments with microglia ablation (53), 1 μl of 7 mg/ml clodronate (HY-B0657A, MCE) or clophosome (F70101C-N, Funakoshi) was injected following the same protocol used for virus injections. As vehicle controls, either ACSF or liposome (F70101-N, Funakoshi) was used. Since no significant difference was observed between the effects of clodronate and clophosome, the results were pooled for analysis.

For behavioral experiments with microglia ablation, 0.5 μl of 3 mg/ml clodronate was injected into the right mPFC (AP +1.1 mm, ML +0.3 mm, DV +2.0 mm).

For drug cannula placement for behavioral experiments, a double cannula (26-gauge, Plastic One or RWD) was placed to target the bilateral mPFC (AP +1.1 mm, ML ±0.3 mm, DV +2.0 mm).

### Acute slice preparation

Acute coronal slices (280 μm thick) containing the mPFC were obtained from 2- to 7- week-old female and male mice. Mice were anesthetized using isoflurane and then perfused transcardially by ice-cold solution (220 mM sucrose, 3 mM KCl, 8 mM MgCl_2_, 1.25 mM NaH_2_PO_4_, 26 mM NaHCO_3_, and 25 mM glucose) and were quickly decapitated. Brains were removed and sliced with a microtome (VT1200, Leica). The slices were incubated at 34°C for 30 minutes and then at room temperature in ACSF (125 mM NaCl, 2.5 mM KCl, 1 mM CaCl_2_, 2 mM MgCl_2_, 1.25 mM NaH_2_PO_4_, 26 mM NaHCO_3_, 20 mM glucose, and 200 µM Trolox (238813, Merck) which was bubbled with 95% O_2_ and 5% CO_2_. Then, the slices were transferred to a recording chamber and perfused with the ACSF solution described above, except using 2 mM CaCl_2_ and 1 mM MgCl_2_, or 2 mM CaCl_2_ and 0 mM MgCl_2_ for magnesium-free experiments. All physiological experiments were performed at 30–32 °C.

### Structural plasticity in slices

Two-photon imaging of dendritic spines was performed using an upright microscope (BX61WI, Olympus) equipped with an FV1200 laser-scanning system (Olympus) with GaAsP photomultiplier tubes (Hamamatsu Photonics) and a water-immersion objective lens (NA 1.0, LUMPlanFI/IR, ×60, Olympus). The system included two mode-locked, femtosecond-pulse Ti:sapphire lasers (MaiTai, Spectra Physics). One was set at a wavelength of 720 nm for the uncaging of glutamate and the other was set at 980 nm for imaging. Fluorescence intensities of green and red fluorophores were detected through 500−550 nm (F03-525/50, Semrock) and 600−680 nm (ET642/80m, Chroma) filters, respectively after a dichroic mirror (T585lpxr, Chroma).

For whole-cell recordings, the patch pipettes (5−8 MΩ) were filled with a solution containing 120 mM potassium gluconate, 20 mM KCl, 10 mM disodium phosphocreatine, 4 mM ATP (magnesium salt), 0.3 mM GTP (sodium salt), 10 mM HEPES, 50 µM Alexa 488 hydrazide (A10436, Thermo Fisher Scientific) or Alexa 594 hydrazide (A10438, Thermo Fisher Scientific) and 5 µM β-actin (APHL99, Cytoskeleton), whose pH and osmolarity were adjusted to 7.25 and 275−280 mOsm by KOH and sucrose, respectively. We did not include Ca^2+^ buffer in the internal solution. For pharmacological experiments, protein kinase A inhibitor (10 µM, 6211, Tocris) was included in the pipette. The cells were voltage-clamped at −70 mV except during stimulation. Cells were discarded if spontaneous action potentials were observed during the stimulating period.

CDNI-glutamate (4 mM, Nard Institute) was locally puffed from a pipette. In experiments with L-noradrenaline hydrochloride (50 µM, #74480, Merck), dopamine hydrochloride (50 µM, D5662, Fujifilm) and clenbuterol (10 μM, C2691, TCI), the reagents were also included in the puff pipette solution. For glutamate uncaging, a 720-nm laser was set to 4−6 mW, depending on the depth to compensate for the decline in stimulation efficiency. Excitatory postsynaptic currents (EPSCs) were recorded by stimulating 3 to 4 spines for 0.6 ms for each while cells were clamped at -70 mV. After confirming the successful induction of EPSCs, structural plasticity was induced by a spike-timing-dependent plasticity (STDP) protocol as previously described (71, 72). Briefly, one burst was composed of three back-propagating action potentials (bAPs, at 100 Hz) preceded by a single EPSP (Δt = +10 ms) induced by two-photon uncaging of CDNI-glutamate. One train of STDP consisted of ten bursts repeated at 10 Hz. Fifteen trains of stimulation were repeated at 0.1 Hz. APs were induced by injection of a depolarizing current of 1.5−2.0 nA for 2 ms. Dendrites containing spines within 100 µm of the soma and 20−50 µm in depth were subjected to plasticity induction. For each experiment, three to four spines were stimulated.

In experiments with D-AP5 (50 µM, #0106, Tocris Bioscience), ICI118,551 (1 μM, ab120808, Abcam), metoprolol (1 μM, M5391, Merck), cilostamide (1 μM, S5806, Selleck), rolipram (1 μM, #180-01411, Fujifilm), deschloroclozapine (DCZ, 100 nM, #7193, Tocris)(*78–80*), TAPI-0 (1 µM, 579050, Calbiochem), pomalidomide (100 μM, P2074, TCI), mouse recombinant tumor necrosis factor-α (TNF-α, 10 nM, 207-13463, Fujifilm), or mouse recombinant interleukin-1β (IL-1β, 1 ng/ml, 094-04681, Fujifilm), slices were constantly perfused with the reagent throughout the series of imaging.

In acute slices following clodronate injection, regions lacking microglia were identified by the absence of fluorescence in transgenic mice expressing tdTomato in microglia. In some experiments, clodronate or vehicle was injected into wild-type mice. Even in these wild-type mice, the ablation site was reliably identified by a needle trace, and cells were recorded within 200 μm of the trace. The site was later confirmed histologically through immunostaining.

### Observational fear learning

Before performing observational fear learning, mice were co-caged for 4 days in their home cage. On day 1, an observer mouse was placed in a conditioning chamber with a stainless-steel rod floor (30 cm × 30 cm, Melquest) for 5 minutes, during which two uncued footshocks (0.4 mA, 2 s each, 1-minute interval; shock generator, No. SG1000-S, Melquest) were delivered. The mice were then returned to their home cage.

On day 2, the mice were placed in a different chamber (17 cm × 35 cm, Melquest), which was divided into two sections: one with a plastic floor for the observer mouse, and the other with a metal rod floor for a naive cagemate (the demonstrator). After 5 minutes of habituation, the demonstrator mouse received 24 successive cued footshocks (tone: 6 kHz, 80−85 dB, 5 s; footshock: 0.7 mA, 2 s, 10-second interval). The observer mouse was then returned to its home cage with its cagemate.

On day 3, memory was assessed by placing the observer mice back into the observation chamber in the same context and presenting three cues (6 kHz tone; 80−85 dB; 60 s; with intervals of 165 s and 177 s). For subthreshold conditioning, 12 cued weak foot shocks (0.2 mA, 2 s each) were presented to the observer mice on day 2.

The stimulus presentation was controlled by triggers generated through a DAQ board (PCIe-6223, National Instruments) using custom-written software. The entire session was recorded with a camera (CS235MU, Thorlab) at 20 fps, synchronized with the stimulus presentation through the DAQ board.

For experiments involving drug infusion, the double cannula was connected to a syringe (1801RN, Hamilton) via a tube containing either the β_2_ blocker ICI118,551 (200 μM in ACSF) or ACSF alone, while the mice were under anesthesia (1% isoflurane, Fujifilm). The drug was infused at a rate of 0.1 μl/min, with a total volume of 1 μl, controlled by a microsyringe pump (Legato 111, KD Scientific). Mice were allowed to recover for at least 30 minutes before undergoing conditioning.

For experiments involving clodronate, mice were euthanized a few days after the memory test, and the patterns of ablation were assessed by immunohistochemistry.

### Auditory conditioning with direct fear experience

On day 1, mice were placed in the conditioning chamber (metal rod floor and plexiglass chamber, stripe wall) for 3 minutes and subsequently received pairs of cues (6kHz tone, 20 s) and footshocks (0.7 mA, 1 s at the offset of tone) for 4 times with 60−79 s pseudo-randomized intervals. On day 2, mice were placed in another context (dot wall) and the cues were presented 4 times with 60−77 s pseudo-randomized intervals. The entire session was recorded with a camera (CS235MU, Thorlab) at 20 fps.

### Thermal place preference test

Thermal preference behavior was assessed using a two-zone apparatus consisting of two adjoining aluminum plates (100 mm × 100 mm each), whose temperatures were independently controlled. The plates formed a continuous flat floor joined by a copper bridge and enclosed by clear acrylic walls (400 mm high), creating a test arena measuring 220 mm × 100 mm. Mice were recorded at 20 frames per second using a side-mounted video camera.

Prior to testing, mice were habituated to the behavioral room for 10 minutes. Each mouse was then placed into the apparatus, with one plate set to a neutral 30 °C and the other to 40 °C. The side assignment of the warmer plate was counterbalanced across animals. Mice were recorded for 10 minutes, and the final 5 minutes were used to calculate place preference (66). Mice that urinated during the session were excluded from analysis.

### Fiber photometry

A 470 nm LED (M470F3, Thorlabs) and 405nm LED (M405F1, Thorlabs) were pulsed (2 ms at 20 Hz, 10 to 15 μW) with shifting 20 ms in order to separate the excitation timing. The excitation light was passed through an excitation filter (ET405/20x; PCR47030x, Chroma Technology), dichroic mirrors (DMLP425, Thorlab; ZT488rdc, Chroma Technology), an objective lens (Olympus, UPlanFLN 20x), then through a rotary joint (RJ-1, Thorlabs) and finally into a low-autofluorescence patch cord (custom made, Thorlabs) connected to the optical implant in the mouse. Emission light was collected through the same patch cord, then passed back through the cube with the dichroic mirror and a fluorescence filter (ET525/50m, Chroma Technology), and then detected by a GaAsP detector (PMT2101/M, Thorlabs). Voltages from the photomultiplier were bandpass filtered (1k Hz) and collected through the DAQ board used for the behavioral experiments.

### Drug administration

For adolescent mice, Tamoxifen (HY-13757A, MedChemExpress, 100 mg/kg body weight) dissolved in corn oil was administered via intraperitoneal injection five times, with 48-hour intervals between doses. Mice were monitored for at least one week before the experiments commenced. For juvenile mice, 20 µg of tamoxifen dissolved in sunflower oil was administered via intragastric injection at P3-5.

### Histology

Mice were anesthetized with isoflurane, perfused using 4% paraformaldehyde (PFA), and decapitated. Brains were post-fixed by 4% PFA overnight and then coronally sectioned at 50 μm using a vibratome (VT-1000, Leica). For immunostaining, slices were incubated in PBS-T (0.2% triton-X in PBS) containing 5% normal goat serum (NGS) at room temperature for 1 h. Slices were then incubated with primary antibodies of rabbit anti-Iba1 antibody (1:1000, #019-19741, Fujifilm) in PBS-T containing 5% NGS for 48 h at 4°C. After washing with PBS-T, slices were incubated with secondary antibodies conjugated with Alexa 488 or Alexa 594 (goat anti-rabbit IgG antibody, 1:500, A-11008 or A-11012, Thermo Fisher Scientific) for 2 h at room temperature. For nuclear staining, TO-PRO-3 (1:2000, T3605, Thermo Fisher Scientific) was incubated simultaneously with the secondary antibodies. The slices were mounted in antifade reagents (H-1000, Vectashield, Vector Laboratories). The slices were subjected to imaging by confocal microscopy (TCS−SP5, Leica) or fluorescent microscopy with structured illumination (BZX-810, Keyence).

Contours were traced by referring to the Allen Mouse Brain Atlas (https://mouse.brain-map.org/static/atlas).

### Data analysis

For spine volume analysis, three-dimensional reconstructions of dendritic morphology were generated by the summation of z-stacked images separated by 0.5 µm. The fluorescence intensity of dendritic spines was normalized by the entire fluorescence of an imaging area. Spine-head volumes were estimated from the total fluorescence intensity. Neighboring spines were defined as spines within 3 µm of the stimulated spines. The baseline value for spine volume was calculated as the mean of the spine volumes at 3 time points before uncaging. To calculate volume change, the baseline was subtracted from the spine volume at each time point. Averaged volume changes at 40 and 50 minutes after STDP stimulation were calculated and were further averaged over spines in the same dendrite for statistical analyses.

For microglia contact analysis, microglial morphology was assessed by summing z-stacked images taken at 1 µm intervals across 5 slices, centered on the plane of maximal intensity for each dendritic spine. At each time point, the presence of microglial processes was recorded if the average signal within a region of interest (ROI) with a 4 µm diameter around the stimulated spine exceeded 3 standard deviations above the background signal of a nearby area.

For image analysis in behavioral experiments assessing the spread of AIP expression or microglial ablation, z-stacked images were processed with a Gaussian filter, and autofluorescence signals from the neocortex were subtracted. Images from all mice were subsequently registered and averaged. For the analysis of AIP expression, the images were binarized prior to averaging. In the AIP experiment for observational fear learning, two mice exhibiting no AIP expression and two mice with the center of AIP expression deviating 0.6 mm anterior to the target were excluded from the analyses. No mice were excluded in the microglial ablation experiment.

For freezing analysis from behavioral data, mouse morphology was detected in each video frame using custom-written scripts with OpenCV in Python. Frame-by-frame differences were then calculated and thresholded to identify freezing responses.

For photometry data, signals from 470 nm and 405 nm excitation were subtracted from autofluorescent reference values. The signal was median-filtered and down-sampled to 2 Hz. Ratio of signal was calculated for each session.

For thermal preference analysis from behavioral data, mouse location was annotated in each video frame using napari in Python. Frame ratios for occupying the lower temperature chamber were then calculated.

### Statistical tests

We used the Kruskal–Wallis test with post hoc Steel’s test or Wilcoxon signed-rank test with Bonferroni correction after Friedman test for multiple comparisons, the Mann–Whitney U test, or the Wilcoxon signed-rank test, depending on the experimental design. Results were considered significant if *P* < 0.05 in two-sided tests. Sample sizes were determined based on previous experiments (71, 72).

---

**Supplementary Figure 1.**
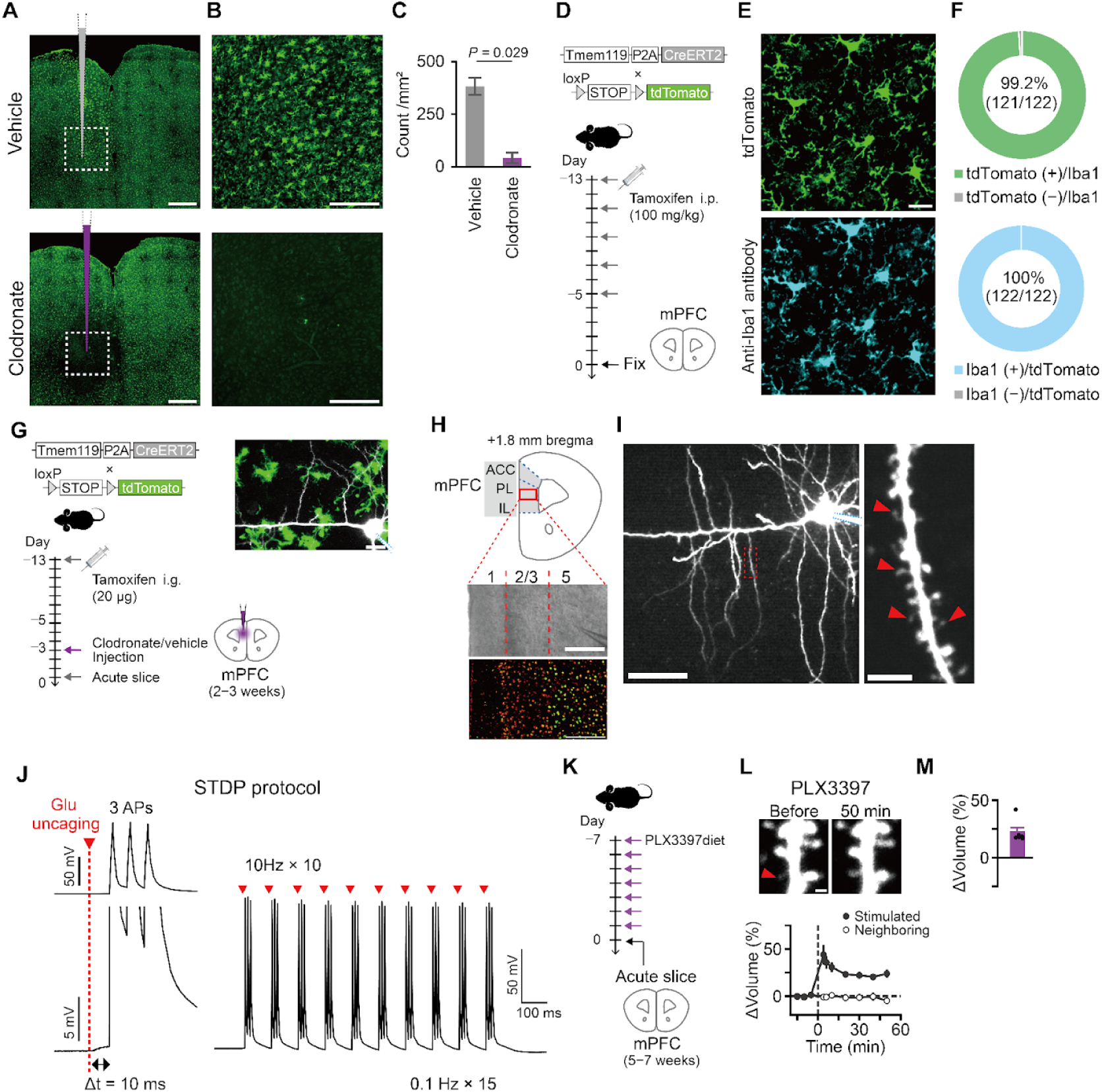
Ablation of microglia and STDP stimulation. **A,B,** Representative images of confocal microscopy showing anti-Iba1 fluorescence in low magnification (A) and high magnification (B) in brains 2 days after vehicle injection (top) and clodronate injection (bottom). Scale bars, 500 μm (A) and 200 μm (B). Dashed lines indicate the areas shown in high magnification. **C,** Plots of microglial counts in the magnified areas shown in (A) and (B) 1–2 days after the injection. n = 4 mice per each condition. Mann–Whitney U test. **D,** Schematic of the experimental timeline in transgenic mice. **E,** Representative confocal microscopy images showing anti-Iba1 and tdTomato fluorescences. Scale bars, 20 μm. **F,** Charts of cell counts from slices of 6 mice. **G,** Schematic of the experimental timeline for microglial ablation in juvenile mice. **H,** Schematics of a coronal slice containing the mPFC (top), and images showing electrode positioning (middle) and immunohistochemical labeling of cortical layers with Satb2 (red) and Ctip2 (green) (bottom). Neurons in the anterior cingulate cortex (ACC), prelimbic cortex (PL), or infralimbic cortex (IL) were recorded. Scale bars, 200 μm. **I,** Images of a pyramidal neuron (left) and dendritic spines (right) obtained by 2-photon microscopy. Red arrows indicate typical spines stimulated by uncaging. Scale bars, 50 μm (left) and 5 μm (right). **J,** Traces of membrane potential during one burst (left) and one train (right) of stimulation in STDP protocols. Action potentials (APs) were evoked by pulsed current injections (1.5−2.0 nA, 2 ms). **K,** Schematic of experimental timeline. adolescent mice were fed PLX3397 containing chow (100 mg/kg of chow) for 7 days. **L,** Representative images of dendrites before and 50 minutes after STDP stimulation (top) and time courses of spine volumes (bottom). Red arrows indicate stimulated spines. Scale bars, 1 μm. **M,** Plots of spine volume changes. Error bars represent S.E.M.

**Supplementary Figure 2.**
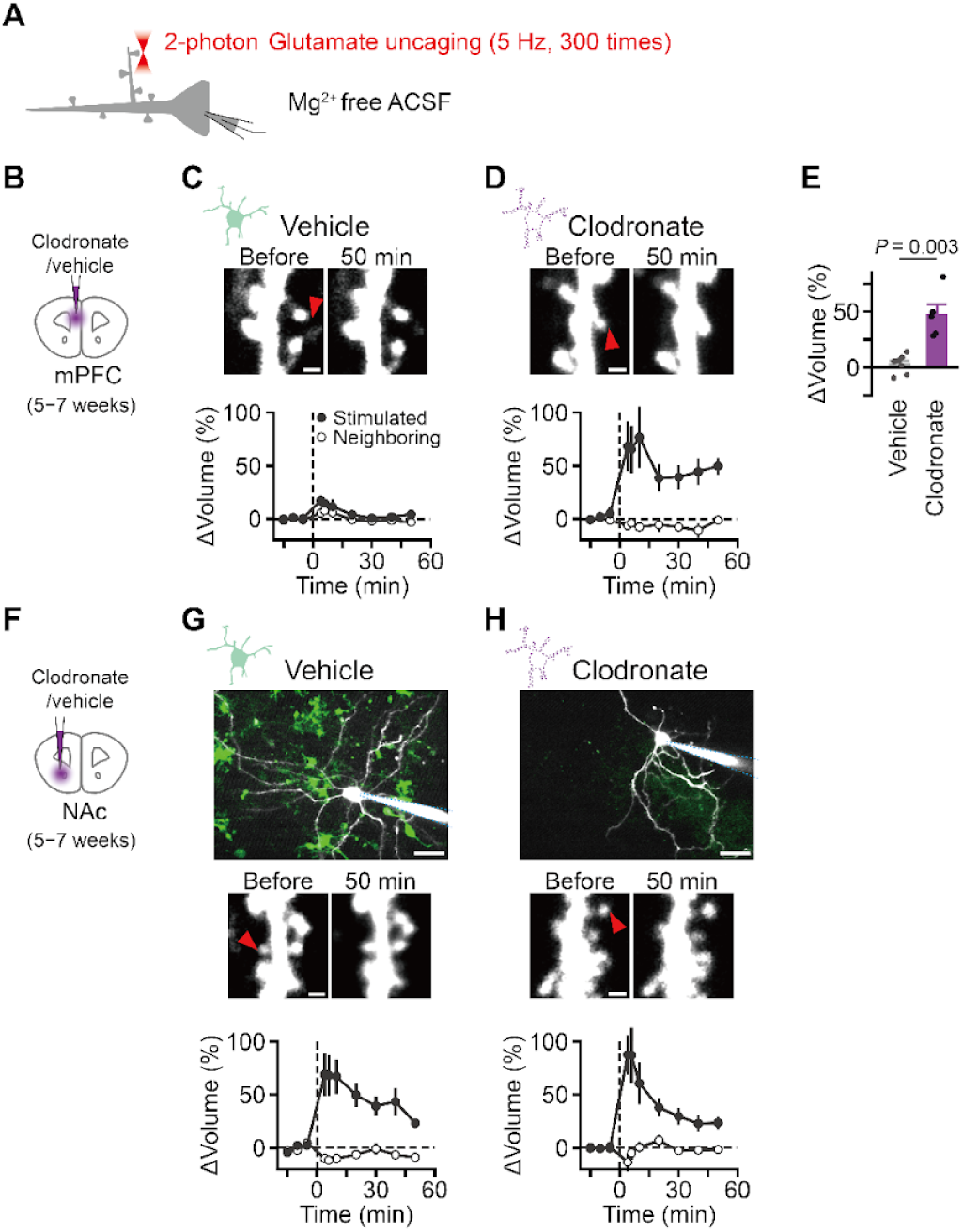
Spine enlargement in magnesium-free condition. **A,** Schematic of the experimental configuration for two-photon uncaging on dendritic spines of a pyramidal neuron under Mg^2+^-free conditions. Whole-cell recordings were made to confirm uncaging-induced EPSCs, but action potentials were omitted during plasticity induction. **B,** Schematic of clodronate or vehicle injection in adolescent mice (5−7 weeks old). **C,D,** Representative images of dendrites before and 50 minutes after STDP stimulation (top) and time courses of spine volumes (bottom) in the slices with vehicle (C) and clodronate (D) injections. Scale bars, 1 μm. **E,** Plots of spine volume changes. **F,** Similar to (B) but injections and experiments were performed in the nucleus accumbens (NAc). **G,H,** Representative images of acute slices in the NAc from Tmem119-CreERT2; Ai14 mice (top) and dendrites before and 50 minutes after STDP stimulation (middle) and time courses of spine volumes (bottom) in the slices with vehicle (G) and clodronate (H) injections. Scale bars, 20 μm (top) and 1 μm (middle). Error bars represent S.E.M. Red arrows indicate stimulated spines. Statistical analysis was performed using the Mann–Whitney U test in (E).

**Supplementary Figure 3.**
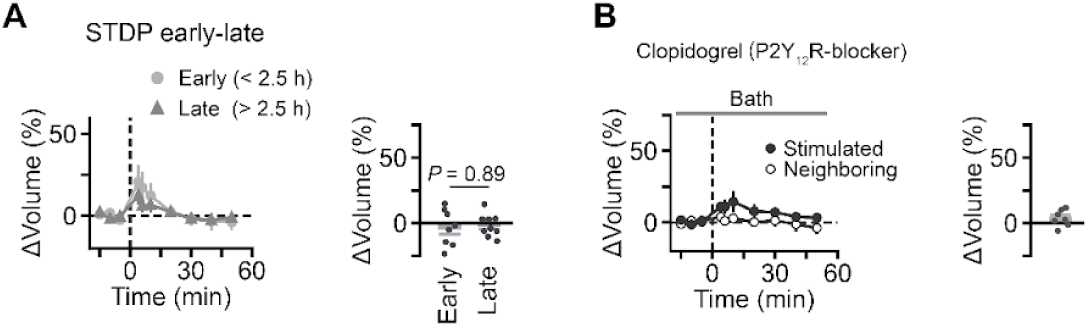
Effects of acute slice preparation. **A,** Time courses of spine volumes before and after 2.5 hours of slice preparation (left) and plots of spine volume changes (right). Data from Figure 1F and Figure 3B were reanalyzed. Incubation times at the time of STDP stimulation were 2.4 ± 0.2 h and 4.5 ± 0.9 h (mean ± SD), respectively. Statistical analysis was performed using the Mann–Whitney U test. **B,** Time courses of spine volumes (left) and plots of spine volume changes (right) in the presence of Clopidogrel (10 μM, P2Y_12_ receptor antagonist). Gray bars indicate the periods for bath application.

**Supplementary Figure 4.**
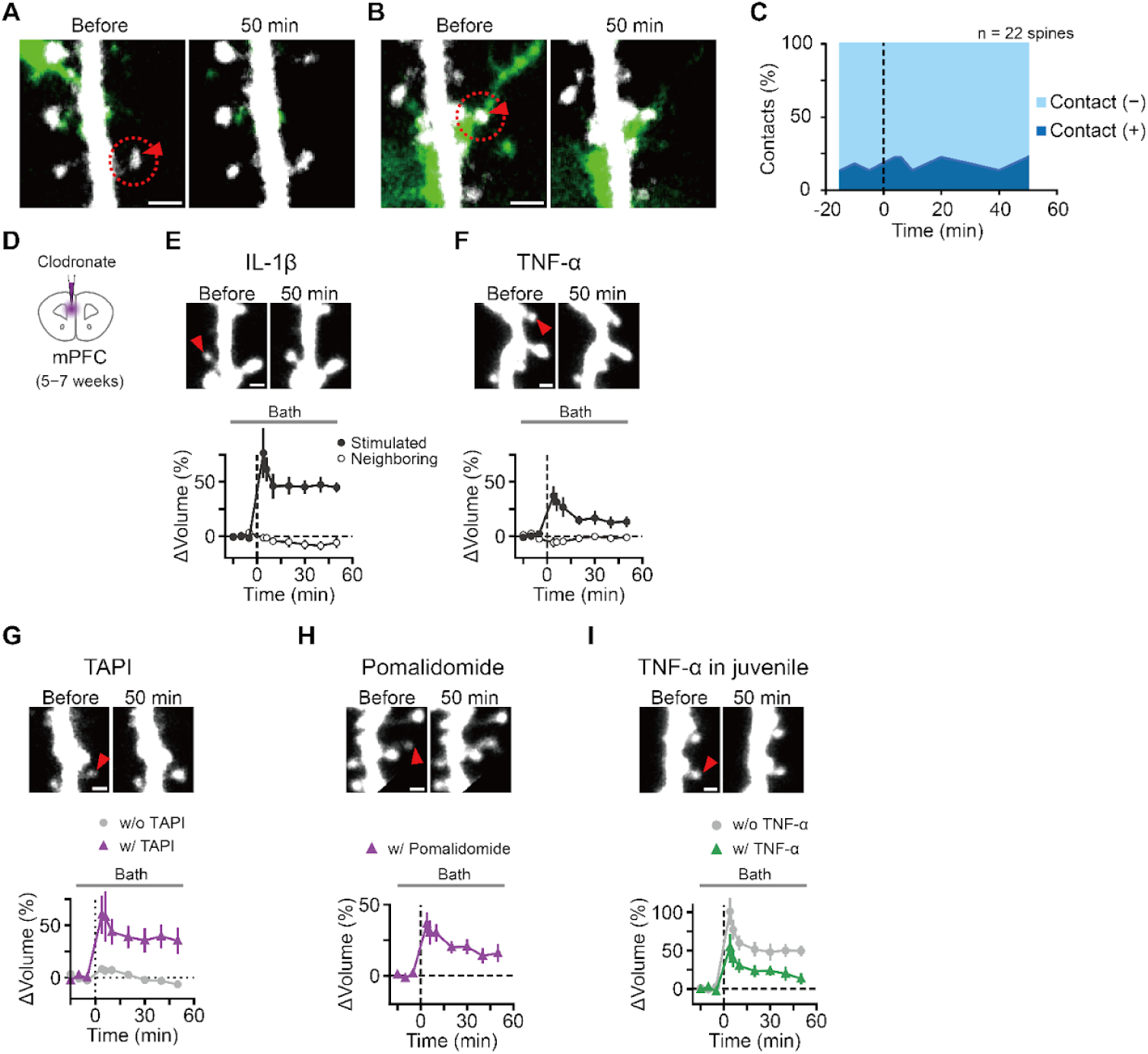
Microglial downstream signaling. **A,** Representative images showing the absence of microglia (green) near the stimulated spine before and after STDP stimulation. The absence of contact was defined by the lack of green signal (within 3 S.D. of a reference background area) in the regions-of-interest (red dashed circle, 4 µm in diameter centered on the stimulated spine) in the 4 µm z-stack image. **B,** Representative images of microglia located near the stimulated spine. **C,** Plots showing the proportion of microglial contacts with stimulated spines over time. **D,** Schematic of clodronate injection in adolescent mice (5−7 week old). **E,F,** Representative images of dendrites before and 50 minutes after stimulation (top) and plots of spine volumes (bottom) in the slices with clodronate injections in the presence of IL-1β (1 ng/ml, bath) (E) and TNF-α (10 nM, bath) (F). **G, H,** Representative images and plots of spine volume changes in the presence of the TAPI-0 (1 µM, bath) (G) and Pomalidomide (100 µM, bath, TNF-α receptor inhibitor) (H) in adolescent mice. **I,** Representative images and plots of spine volume changes in the presence of TNF-α (10 nM, bath) in juvenile mice (2−3 weeks old). Red arrows indicate stimulated spines. Scale bars, 2 µm in (A, B), 1 µm in (E, F, G, H, I). Error bars represent S.E.M.

**Supplementary Figure 5.**
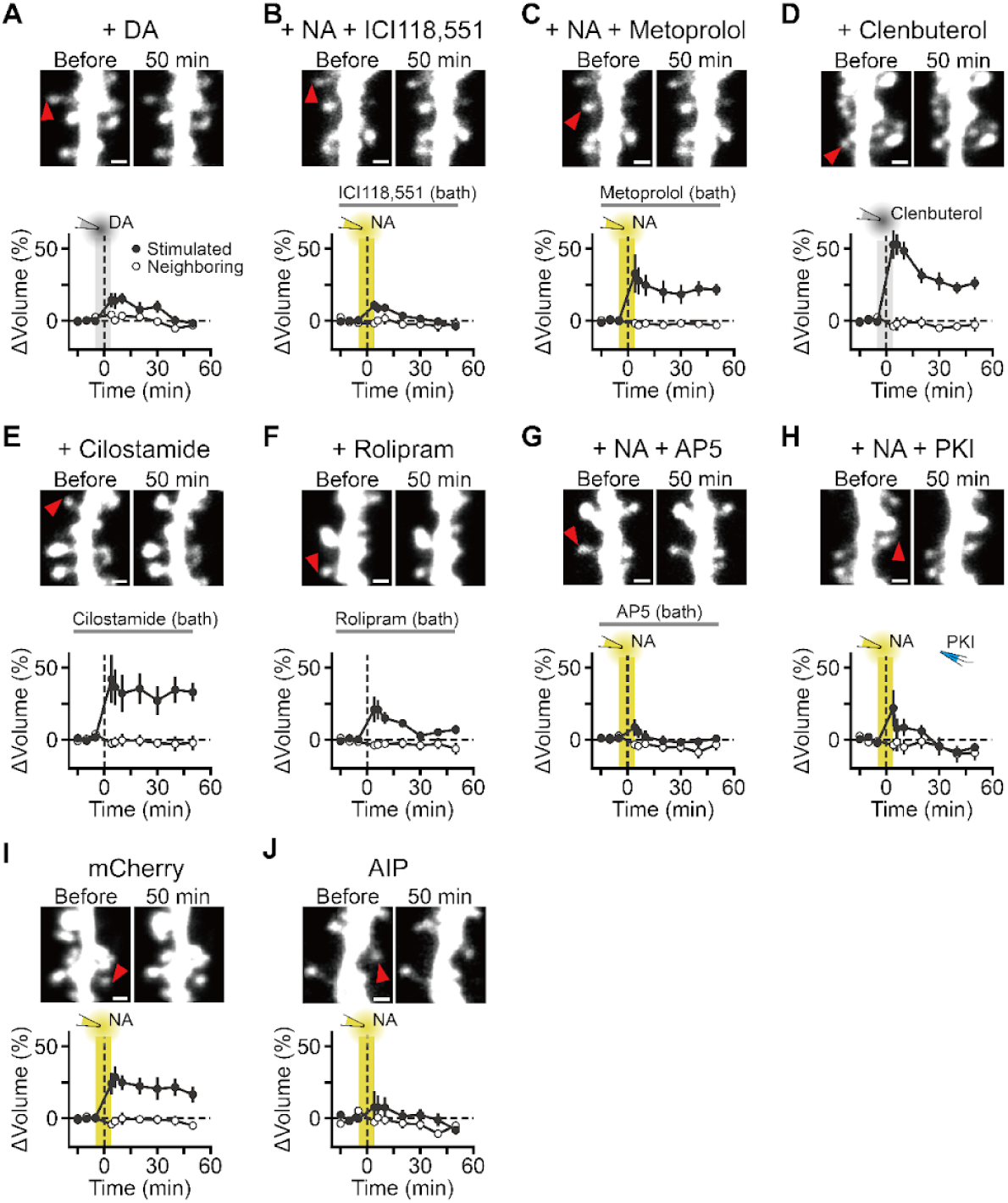
Pharmacological studies on spine enlargement in the mPFC. **A–H,** Representative images of dendrites before and 50 minutes after stimulation (top) and time courses of spine volumes after STDP stimulation (bottom) in the presence of (A) dopamine (DA), (B) NA and a β_2_ blocker, (C) NA and a β_1_ blocker, (D) a β_2_ agonist, (E) PDE3 inhibitor, (F) PDE4 inhibitor, (G) NA and AP5, and (H) NA and PKI. Shading indicates the periods of puff application for DA, NA or a β_2_ agonist. Gray bars indicate the periods for bath application. Error bars represent standard error. Filled and open symbols represent spine volumes measured from stimulated spines and unstimulated neighboring spines (control), respectively. Red arrows indicate stimulated spines. **I, J,** Representative images of dendrites before and 50 minutes after stimulation (top) and time courses of spine volumes (bottom) in the absence (I) and presence (J) of AIP. Scale bars, 1 µm. Error bars represent S.E.M.

**Supplementary Figure 6.**
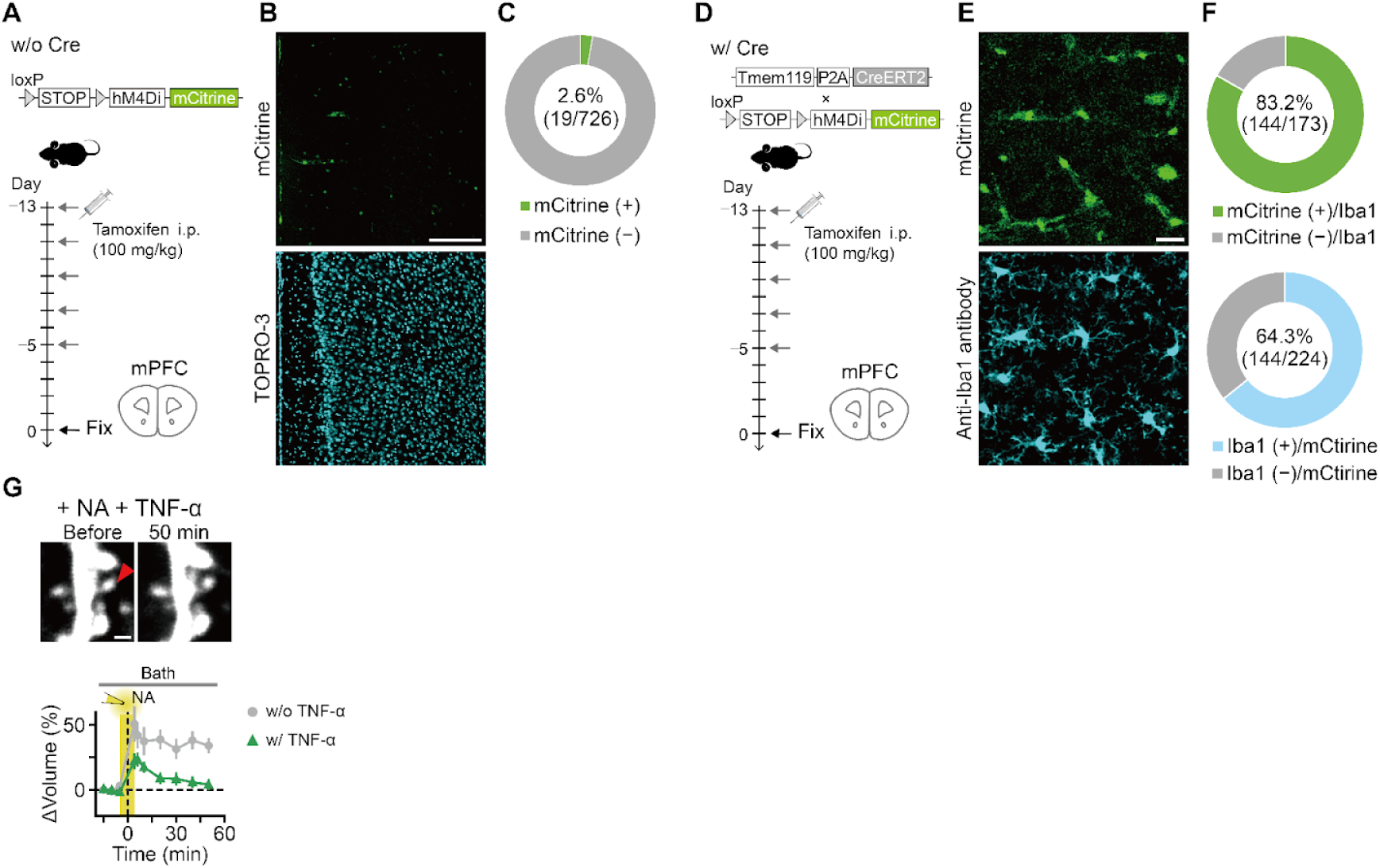
Chemogenetic manipulation of microglia. **A,** Schematic of the experimental timeline in mice without the Cre-driver. **B,** Images showing mCitrine fluorescence (top) and nuclear staining with TOPRO-3 (bottom) in mice without the Cre-driver. **C,** Chart showing proportion of mCitrine expressing cells out of TOPRO-3 positive cells. Data were obtained from slices of 2 mice. **D,** Schematic of experimental timeline in mice with Cre-driver. **E,** Images showing mCitrine fluorescence (top) and immunostaining with anti-Iba1 antibody (bottom) in mice with Cre-driver. **F,** Chart showing proportion of the mCitrine expressing cells out of anti-Iba1 positive cells (top) and anti-Iba1 positive cells out of mCitrine expressing cells (bottom). Slices from 6 mice. **G,** Representative images of dendrites before and 50 minutes after STDP stimulation (top) and time courses of spine volumes (bottom) with or without TNF-α (10 nM, bath) in the presence of NA (50 µM). Red arrows indicate stimulated spines. Scale bars, 200 μm in (B), 20 μm in (E), and 1 μm in (G). Error bars represent S.E.M.

**Supplementary Figure 7.**
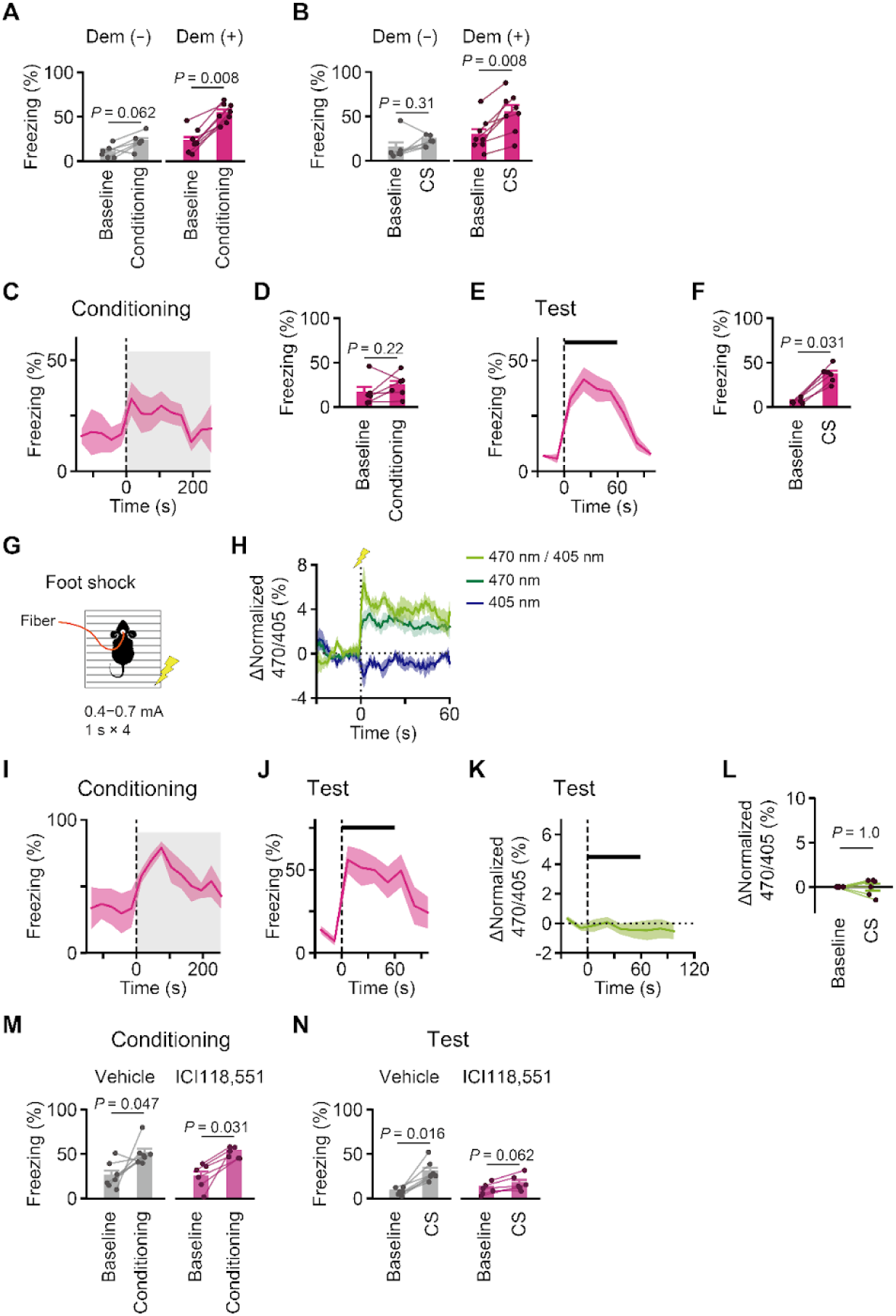
Observational fear learning. **A, B,** Plots of freezing on day 2 of conditioning (A) and day 3 of test (B) of mice conditioned with or without demonstrator. **C−F,** Traces (C,E) and plots (D,F) of freezing on day 2 of conditioning (C,D) and day 3 of test (E,F) in female mice. A pair of female mice was used for observer and demonstrator. **G,** Schematic of photometry recording in response to foot shock (0.4−0.7mA, 1 s). **H,** Traces of raw fluorescences (signals obtained by excitation with 405 nm LED or 470 nm LED) and ratio (470 nm/405 nm) against the onset of the foot shock. Averages of 4 trials are shown. Note that the isosbestic wavelength of the probe is around 410 nm and therefore the signal with 405 nm excitation exhibited the opposite direction to that of 470 nm excitation. These mice were a separate set from those used in observational fear learning. **I,J,** Traces of freezing on day 2 (I) and day 3 (J) in fiber photometry recording experiment. **K,L,** Trace (K) and plots (L) of GRAB-NE_2m_ fluorescence on day 3. **M,N,** Plots of freezing on day 2 (M) and on day 3 (N) in ICI118,551 experiments. Wilcoxon signed-rank test for (A, B, D, F, L, M, N). Error bars and shades indicate S.E.M.

**Supplementary Figure 8.**
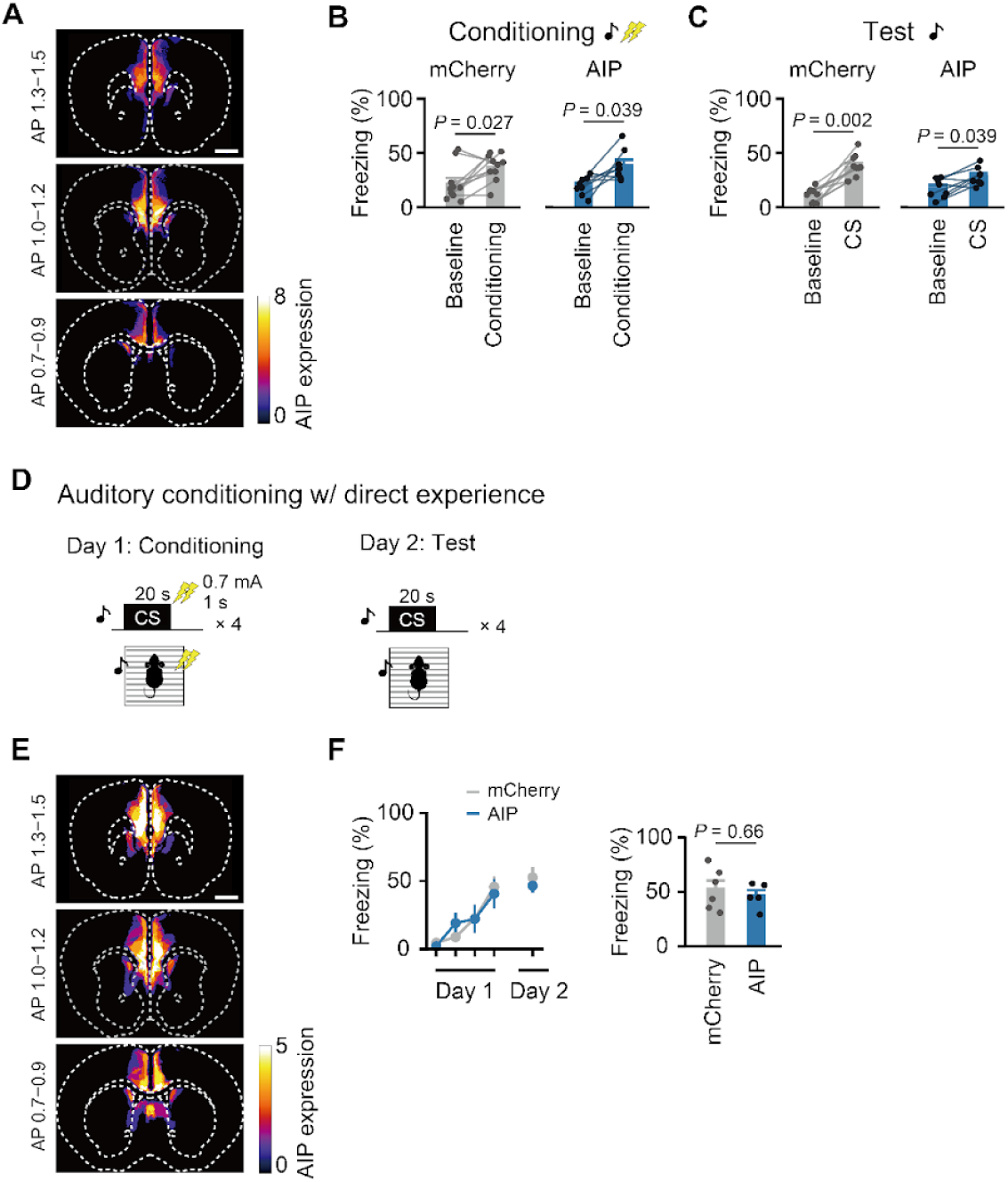
Inhibition of CaMKII signaling in fear learning. **A,** Images showing averaged intensity of AIP expression in all the mice engaged in the observational fear learning. AP indicates anteroposterior coordinates from the bregma in mm. **B, C,** Plots of freezing on day 2 (B) and on day 3 (C) in AIP experiments. **D,** Schematic of auditory conditioning with direct fear experience. **E,** Images showing averaged intensity of AIP expression in all the mice engaged in auditory conditioning with direct fear experience. **F,** Time courses of freezing on day 1 (conditioning) and on day 2 (test) (left) and plots on day 2 (right). Freezing responses during tone periods (20 s) are shown for each trial (day 1) or average of 4 trials (day 2). Scale bars in (A, E) indicate 1 mm. Wilcoxon signed-rank test for (B,C), Mann–Whitney U test for (F).

**Supplementary Figure 9.**
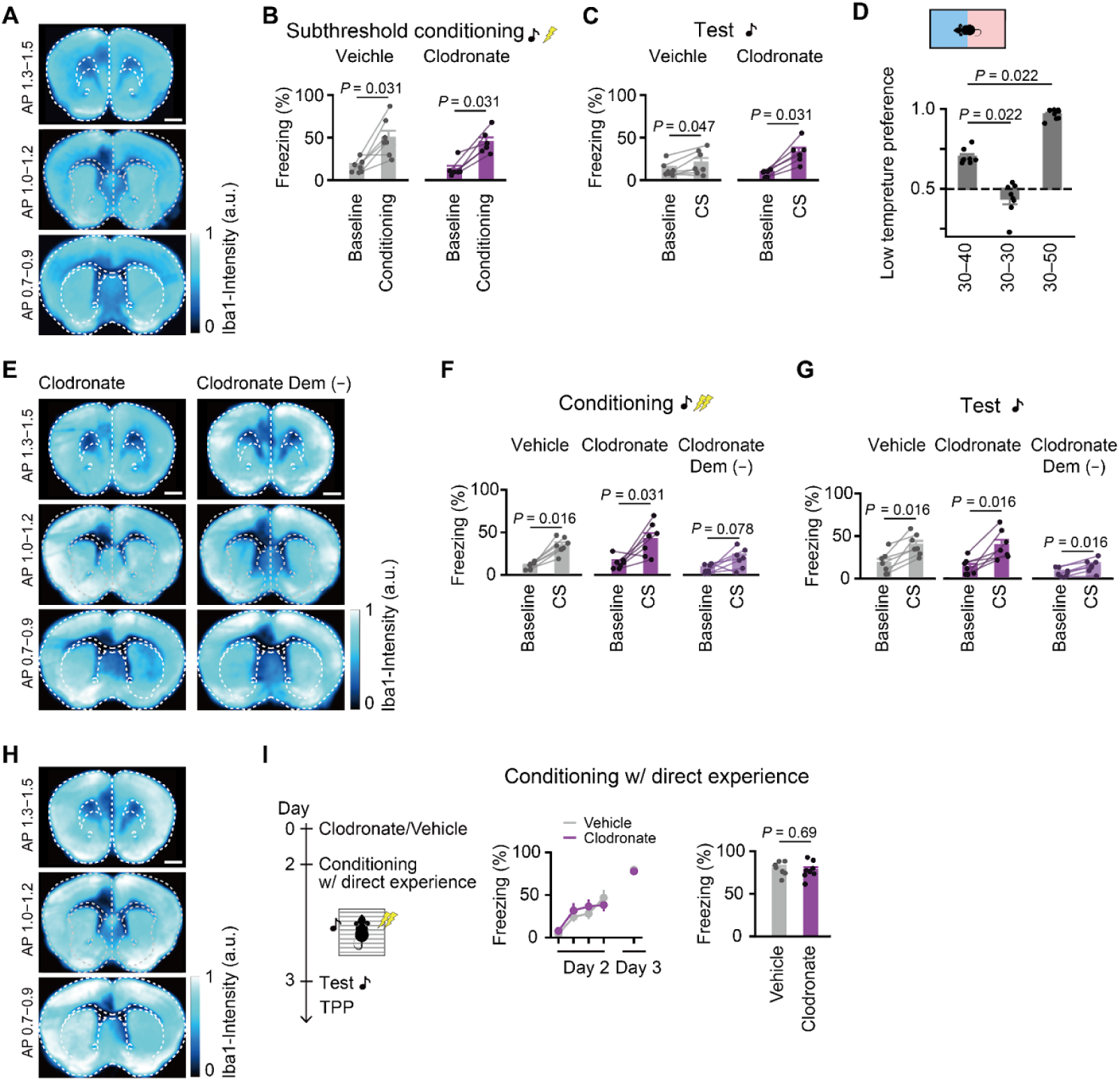
Ablation of microglia in fear learning. **A,** Images showing averaged intensity of Iba1 immunoreactivity in the brain from all the clodronate-injected mice used for subthreshold conditioning. **B,C,** Plots of freezing on day 2 (B) and on day 3 (C) in subthreshold conditioning. **D,** Thermal place preference with various temperatures. Times spent in the lower temperature plate during the latter 5 min out of total 10 min recording periods are shown. **E,** Images of Iba1 immunoreactivity from all the mice used for conditioning and thermal place preference test. **F, G,** Plots of freezing on day 2 (F) and on day 3 (G) in conditioning with clodronate injections with and without demonstrator. **H,** Images of Iba1 immunoreactivity from all the mice used for conditioning with direct experience. **I,** Auditory fear conditioning with direct experience in the presence and absence of clodronate. Scale bars in (A, E, H) indicate 1 mm. Wilcoxon signed-rank test for (B,C, F, G), Wilcoxon signed-rank test with Bonferroni correction after Friedman test (*P* < 0.001) for (D), Mann–Whitney U test for (I).

**Supplementary Figure 10.**
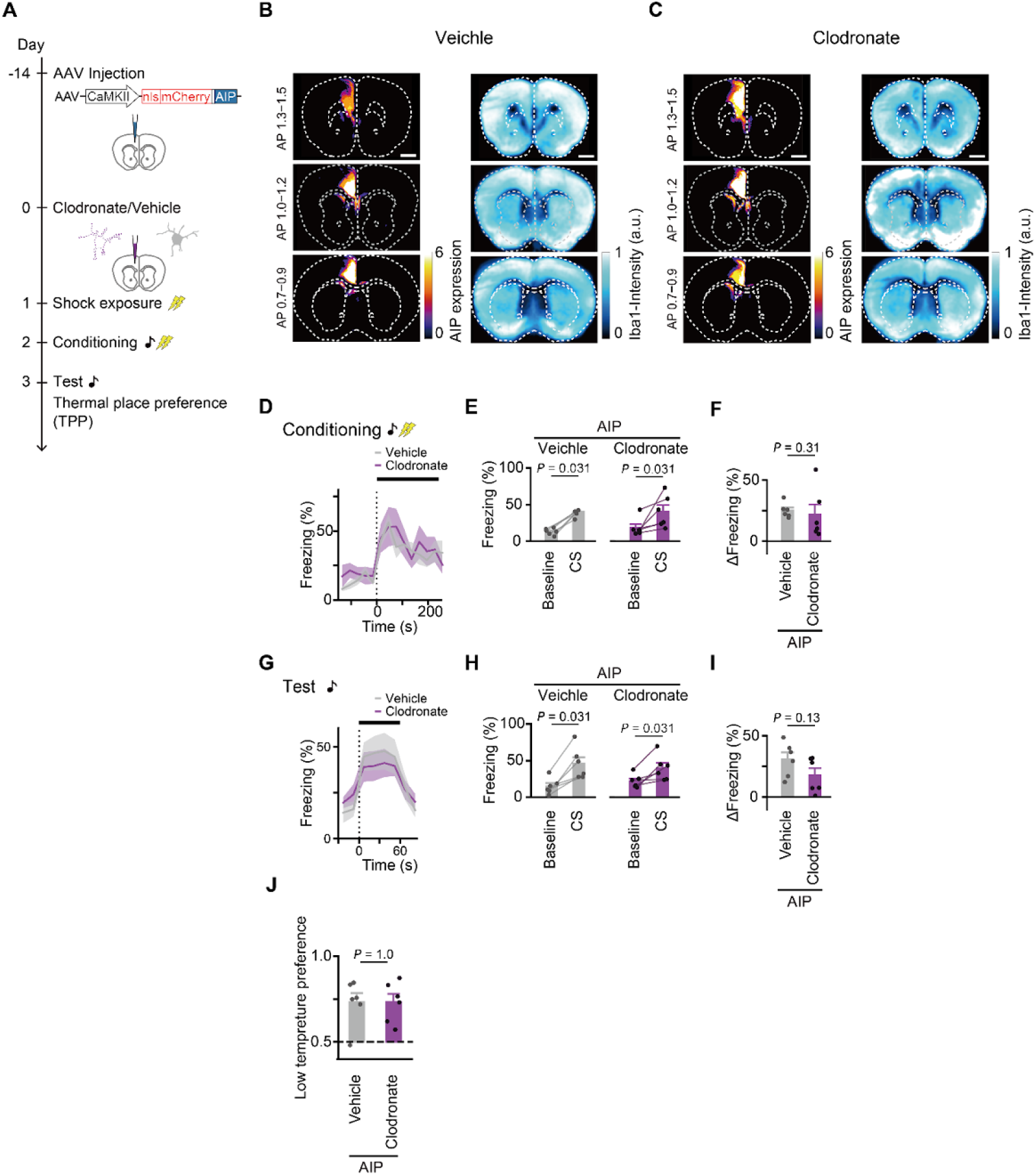
Occlusion of microglial ablation effects on thermal sensitivity. **A,** Schematic diagram for experimental timeline. The procedure is the same as in Figure 5F, except that mice were injected with AIP-expressing AAV in the right mPFC. **B,C,** Images showing averaged AIP expression intensity (left) and Iba1 immunoreactivity (right) in the brain from all mice engaged in the task with vehicle (B) and clodronate (C) injections. **D−I,** Traces and plots of freezing on day 2 (D, E, F) and on day 3 (G, H, I). **J,** Plots of thermal place preference. Wilcoxon signed-rank test for (E, H), Mann–Whitney U test for (F, I, J).

## Notes

### Competing Interest Statement

The authors have declared no competing interest.

### Summary of Updates

We revised the title, abstract, introduction, and discussion.

